# Phylogenomics resolves the relationship and the evolutionary history of planthoppers (Insecta: Hemiptera: Fulgoromorpha)

**DOI:** 10.1101/2024.07.26.605304

**Authors:** Junchen Deng, Adam Stroiński, Jacek Szwedo, Hamid Reza Ghanavi, Etka Yapar, Diego Castillo Franco, Monika Prus-Frankowska, Anna Michalik, Niklas Wahlberg, Piotr Łukasik

## Abstract

Planthoppers (Hemiptera: Fulgoromorpha) are a species-rich and globally distributed insect clade with high economic, ecological, and evolutionary importance. However, the relationships among planthopper lineages and families remain unclear. Previous efforts based on inconsistent morphological traits, a few genes, or limited sampling often resulted in conflicting tree topologies. Here, we used genome-level data to assemble 1164 nuclear single-copy genes and 13 mitochondrial protein-coding genes for 149 planthopper species representing 19 out of 21 extant families. Additional markers were added from published mitogenomes, expanding our sampling to 285 species. These markers were used for Maximum Likelihood-based tree inference and dating analyses. The newly inferred phylogenies validated well-accepted relationships and recovered novel placements. Taxonomic conclusions include the establishment of a new family Borysthenidae **stat. rev.** within Delphacoidea and a new superfamily Meenoploidea **superfam. nov.** including redefined Kinnaridae **stat. rev.** and Meenoplidae **stat. rev.**, the confirmation of the monophyletic family Achilixiidae outside the Achilidae-Derbidae clade, and the transfer of tribes Lyncidini and Amyclini to Dictyopharidae and the genus *Madagascaritia* to Fulgoridae. The time analyses based on 57 nuclear markers and 30 fossils dated the origin of crown Fulgoromorpha back to Guadalupian, Permian (∼263 Ma), close to the maximum constraint at 267.3 Ma, while applying an older root constraint resulted in an origin in Mississippian, Carboniferous (∼332 Ma). While future sampling of unstudied fauna in unexplored regions or habitats may change the topology, the current phylogenomic analysis will serve as a solid foundation for research into planthopper ecology, evolution, and significance.

## Introduction

Planthoppers (Hemiptera: Fulgoromorpha) are a clade (suborder or infraorder, depending on the authority; see Bartlett et al., 2018; Szwedo, 2017) of sap-feeding insects comprising 21 extant families, 16 extinct families, and >14000 named species. They are distributed worldwide, except for mainland Antarctica, occupying diverse ecosystems from tropical, subtropical, and temperate forests through grasslands, to deserts (Bourgoin, 2024). Many species are considered economically significant as crop pests through direct feeding damage, vectoring of plant pathogens, and the excretion of honeydew that encourages mold growth and can impede photosynthesis (Dietrich, 2009; Heong et al., 2015). Through herbivory, planthoppers also play significant roles in natural ecosystems. At the same time, they serve as an important food resource for other organisms, including ants, bees, geckos, and even local people, through predation or honeydew production (Bourgoin et al., 2023; Dietrich, 2009; Urban and Cryan, 2007). In addition to ecological importance, they are of cultural interest. In particular, the variation in head shape and wing venation makes many species, especially the strikingly large and colored ones from the family Fulgoridae, popular to taxonomists and nature lovers (O’Brien, 2002; Urban and Cryan, 2009). Moreover, planthoppers can teach us about more general ecological and evolutionary processes and patterns through their symbiotic relationships with diverse bacteria and fungi (Bennett and Moran, 2013; Deng et al., 2023; Michalik et al., 2023). The evolution happening in the genomes of their ancient symbionts has paralleled with the genomes of organelles, providing insights into such ancient events as the origin of eukaryotes (McCutcheon et al., 2019).

To better understand the evolution of morphological traits, life histories, ecological functions, and microbial symbioses across clades and species, a robust phylogeny of worldwide planthoppers is needed. However, relationships among planthopper families are still under debate. Early efforts to establish relationships between families date back to the 20^th^ century when molecular tools were not commonly used and researchers tended to rely on morphological and ecological traits to infer species phylogenies (Bourgoin, 1993; Emeljanov, 1990; Liang and Yang, 2001; Muir, 1923). However, there was little congruence among the various topologies proposed. In the late 20^th^ century and early 21^st^ century, with molecular approaches slowly gaining ground in taxonomic work, a few studies attempted to reconstruct the phylogeny of planthopper families using a single molecular marker (e.g., ribosomal DNA) (Bourgoin et al., 1997; Yeh et al., 2005). Despite the poor resolution, these studies agreed on a few patterns, such as Delphacidae, Cixiidae, and Meenoplidae being among the most anciently diversifying lineages while Tettigometridae is younger, and Dictyopharidae and Fulgoridae being sister taxa.

As sequencing technology advances, several studies used more markers to investigate the relationship between planthopper families. Urban and Cryan (2007) performed the first multi-gene analysis on 71 species from 19 families using four nuclear genes (18S rDNA, 28S rDNA, Histone subunit 3, and Wingless) with a total alignment length of ∼4.3 kbp. Later on, in 2012, they performed a similar analysis on 77 species from 18 families by adding one more mitochondrial marker (cytochrome oxidase I) to the nuclear dataset (Urban and Cryan, 2012). Afterward, Song and Liang (2013) increased the sampling size to 133 species for the marker 18S rDNA and performed analyses on either a single marker or the combined dataset of two nuclear markers (18S rDNA and 28S rDNA) and two mitochondrial markers (16S rDNA and cytochrome b). More recently, Bucher et al. (2023) published a phylogeny of 531 species from 19 families based on three nuclear markers (18S rDNA, 28S rDNA, and Wingless) and two mitochondrial markers (cytochrome oxidase I and cytochrome b). In the same year, Wang et al. (2023) published a phylogeny based on the complete mitogenomes (13 protein-coding genes and two rDNAs) of 113 species from 17 families. These studies recovered several sister groups in common, including lineages Delphacidae-Cixiidae, Kinnaridae-Meenoplidae, Fulgoridae-Dictyopharidae, and Lophopidae-Eurybrachidae, and indicated that lineages Delphacidae-Cixiidae and Kinnaridae-Meenoplidae diverged early from the remaining planthoppers. However, the relative placements of other clades are still uncertain, and the relationships among the remaining families are far from clear. Certain patterns were recovered by some studies but not the others, such as the monophyly of the Achilidae-Achilixiidae-Derbidae clade in Urban and Cryan (2007) and Bucher et al. (2023), the Tettigometridae-Caliscelidae in Song and Liang (2013), and the Flatidae-Ricaniidae clades in Song and Liang (2013) and Wang et al. (2023). On the other hand, the usefulness of mitochondrial genomes for resolving deep phylogenetic relationships has been questioned in other groups of insects (Ghanavi et al., 2022). These uncertainties call for comprehensive studies and analyses with more genetic markers using more advanced methods.

In the last decade, the fast advance of high-throughput sequencing technology has enabled the sequencing of thousands of gene markers at a reasonably low cost, ushering in the era of phylogenomics. Two important studies addressing phylogenetic questions at higher taxonomic levels (e.g., order and suborder) have confirmed the status of crown Fulgoromorpha (Delphacoidea and Fulgoroidea) as a monophyletic clade that is sister to crown Cicadomorpha: Clypeata (cicadas, spittlebugs, and leafhoppers/treehoppers), with the two clades often grouped into a suborder Auchenorrhyncha (Bartlett et al., 2018; Davranoglou and Hartung, 2024; Szwedo, 2017). On the other hand, they provided limited information on the relationships among planthopper clades. Johnson et al. (2018) used transcriptome data to reconstruct the history of diversification of hemipteroid insects, including 13 planthopper species from nine families, suggesting that Fulgoromorpha separated from Cicadomorpha ∼300 million years ago (Ma). Later, Skinner et al. (2020) inferred the phylogeny of all extant auchenorrhynchan superfamilies using a transcriptomic dataset that includes 25 planthopper species from 15 families. Their phylogenetic analyses, based on >2000 orthologs, provided strong support for some sister groups, including Delphacidae-Cixiidae, Fulgoridae-Dictyopharidae, Tettigometridae-Caliscelidae, and Acanaloniidae-Flatidae but the relationships among the rest of families remain unclear. Phylogenomic efforts spanning comprehensive global collections of planthoppers from major clades are still missing.

Here, we attempted to reconstruct the evolutionary history of worldwide planthoppers through phylogenomic inference based on high-throughput genomic sequencing. To resolve the phylogenetic relationships, we prepared datasets including 1164 single-copy nuclear genes assembled from metagenomes of 149 planthopper species spanning 19 out of 21 recognized extant families. In addition, we enlarged the nuclear dataset with complete mitochondrial genomes extracted from GenBank and our metagenomic assemblies, reaching a total of 285 planthopper species. To estimate the date of origin of major clades, we carefully curated 30 fossils as calibration points on our best phylogeny of 149 species. Moreover, we tested how different sets of fossil priors and age constraints affect our results.

## Materials and Methods

### Sample processing and metagenomic sequencing

Individuals of 151 planthopper species from 19 families were sampled ethically following all applicable international and local permitting requirements from their natural habitats around the world (Table S1). The remaining two established families, Gengidae and Hypochthonellidae, known only from limited specimens collected in southern Africa and, to our knowledge, with no molecular data published, are not included due to sampling difficulties. For all individuals, we extracted DNA from either the abdomen or bacteriome – an organ hosting symbiotic bacteria, using various DNA extraction kits (Table S1). Metagenomic libraries of each individual were prepared with NEBNext Ultra II DNA Library Prep Kit (BioLabs, New England) and sequenced on Illumina HiSeq 2500 in a Rapid Run mode (2x250bp), HiSeq X, or NovaSeq 6000 S4 (2x150bp reads) (Table S1).

### Single-copy nuclear gene assembly and multiple sequence alignment

We prepared the multiple sequence alignment (MSA) for phylogenetic analysis in the following steps: 1) the preparation of a custom single-copy gene reference database; 2) single-copy gene assembly from metagenomic data; 3) alignment and filtering. The analysis pipeline and the custom Python scripts can be found on the GitHub page (https://github.com/junchen-deng/phylogenomic_analysis_pipeline).

To enable read mapping and gene assembly from generally low-coverage genomic datasets, we first assembled our custom single-copy gene database for planthoppers using BUSCO v5.4.4 (Manni et al., 2021). The hemipteran ortholog database (hemiptera_odb10, v2020-08-05), including 2510 single-copy orthologs, was downloaded from BUSCO. Then, we extracted from GenBank genome assemblies of species covering a wide range of evolutionary distances to planthoppers. This included chromosome-level assemblies from pea aphid *Acyrthosiphon pisum* (Sternorrhyncha: Aphididae), glassy-winged sharpshooter *Homalodisca vitripennis* (Cicadomorpha: Clypeata: Membracoidea), sugarcane spittlebug *Callitettix versicolor* (Cicadomorpha: Clypeata: Cercopidae) and brown planthopper *Nilaparvata lugens* (Fulgoromorpha: Delphacoidea), and scaffold-level assembly from ‘*Zanna*’ *intricata* (synonym to *Pyrops intricatus*; Fulgoromorpha: Fulgoroidea) (Table S1). We then fed these assemblies to BUSCO and identified 2447, 2098, 2340, 2322, and 1168 complete single-copy genes, respectively. Finally, we combined these five datasets with busco_ancestral from the BUSCO hemiptera_odb10 dataset and generated the custom database – in amino acid – for the next step.

In the second step, we assembled the single-copy genes (orthologs) presented in the custom database using the metagenome of each species. We first cleaned Illumina paired-end reads in each metagenomic dataset by adapter trimming and quality control using trim_galore v0.6.4 (settings: --length 80 -q 30; https://github.com/FelixKrueger/TrimGalore). The quality of clean reads was confirmed by FastQC v0.11.9 (https://github.com/s-andrews/FastQC). Then, HybPiper v2.1.2 was used to assemble single-copy genes in each clean metagenomic dataset (Johnson et al., 2016). HybPiper first performed target enrichment by blasting and mapping Illumina reads against the custom database using DIAMOND v2.0.15 (setting: -t_aa custom_db --diamond --diamond_sensitivity sensitive) (Buchfink et al., 2021). The clustered reads for each single-copy gene were then assembled with SPAdes v3.15.5 implemented in Hybpiper (Prjibelski et al., 2020). Due to the low coverage of the host genome in our metagenomic dataset, we reduced the coverage cutoff for SPAdes assembly to four (setting: --cov_cutoff 4) to increase the contig length without losing much base-level accuracy. This allowed us to assemble all 2510 single-copy genes presented in the custom database from at least some samples.

In the final step, we implemented several rounds of filtering to obtain clean alignments for each ortholog. We first ran hybpiper paralog_retriever to detect and remove potential paralogs. This reduced the dataset to 2313 genes. Then, we ran Fast Statistical Alignment (FSA) with fsa v1.15.9 to generate alignments of amino acids for each gene including the target species and two outgroups (*H. vitripennis* and *C. versicolor*) (Table S1; Bradley et al., 2009); FSA allows more reliable identification of non-homologous sequence, which improves overall alignment quality in our case because our data contained some reads of other non-target eukaryotic sources (e.g., symbiotic fungi and insects other than planthoppers, such as putative parasitoids) in our metagenomic data when using a low coverage cutoff for spades assembly in the previous step. Next, trimAl v1.4 was used to trim the amino acid alignments (setting: -automated1) (Capella-Gutierrez et al., 2009). The nucleotide alignments were generated by reverse translating the trimmed amino acid alignments based on the column numbers output by trimAl v1.4 (setting: -colnumbering). Then, we ran other custom Python scripts to filter trimmed alignments by keeping genes that 1) are longer than 200 amino acids and 2) were reconstructed for at least 75% of all target species (i.e. 112 out of 149 species in our case) with sequence length above 25% of the gene length. This further reduced the dataset to 1508 genes. Finally, we ran FastTree v2.1.11 to generate a Maximum Likelihood tree for each gene (setting: -nt -gtr) (Price et al., 2010). We then manually examined each gene tree by looking for unusual topologies (e.g., long branches of a single, few, or a clade of species); long branches of a single or few species can result from short sequence length or contamination while long branches of a clade of species may indicate potential paralogs not detected by Hybpiper. The alignments of gene trees with unusual topologies were further examined manually. The problematic alignments were either cleaned by manually cutting out the troublesome regions or sequences or excluded from the subsequent analyses. At this stage, we also decided to exclude two species, *Stenocranus major* and *Parahydriena hyalina acuta*, which were placed repeatedly with outgroups due to significant metagenome contamination from the DNA of putative parasitoid insects. In the end, we reached a final list of 1164 single-copy orthologs from 149 species.

### Mitogenome assembly, annotation, and mitochondrial marker extraction

The clean Illumina reads obtained in previous steps after adaptor trimming and quality control were assembled using MEGAHIT v1.1.3 with k-mer sizes from 99 to 255 (Li et al., 2016). We then used NanoTax.py (https://github.com/junchen-deng/NanoTax), which performs blast searches using assembled contigs against a custom database containing previously published mitogenomes, to identify mitochondrial contigs. Complete or partial mitogenomes of 149 out of 151 planthopper species were assembled from metagenomes. The mitogenomes of *S. major* and *P. hyalina acuta* from contaminated metagenomes were successfully assembled and verified while the mitogenomes of two other species, *Bebaiotes* sp. and *Omolicna uhleri*, failed to assemble (Table S1). The identified mitochondrial contigs were then annotated with a custom Python script modified from (Łukasik et al., 2019). The script first extracted all the Open Reading Frames (ORFs) and their amino acid sequences from a mitogenome. Then, the ORFs were searched recursively using HMMER v3.3.1 (Eddy, 2011), through custom databases containing manually curated sets of planthopper mitochondrial protein-coding and rRNA genes. The rRNA genes were searched with nhmmer implemented in HMMER v3.3.1 (Wheeler and Eddy, 2013), and tRNAs were identified with tRNAscan-SE v2.0.7 (Chan et al., 2021). All 13 mitochondrial protein-coding genes (PCGs) were annotated and extracted for each species. Additionally, we extracted mitochondrial markers from previously published mitogenomes of 134 planthopper species on GenBank (published before February 2024; Table S1), mostly from Eastern Palearctic and Oriental taxa not well-represented in the primary dataset. Mitogenomes of four outgroups from Cicadomorpha, including the keeled treehopper *Entylia carinata* (superfamily Membracoidea), the cicada *Tettigades undata* (superfamily Cicadoidea), the meadow spittlebug *Philaenus spumarius* (superfamily Cercopoidea), and the leafhopper *Macrosteles quadrimaculatus* (superfamily Membracoidea), were included in the annotation (Table S1). The amino acid alignment of each mitochondrial marker was inferred with Mafft v7.475 (settings: --localpair --maxiterate 1000; Katoh and Standley, 2013). The nucleotide alignments were generated by reverse translation of amino acid alignments as described in the previous steps.

### Phylogenetic inference

We prepared four concatenated datasets, with both amino acid and nucleotide alignments included in each dataset, for the phylogenetic analyses. These were 1) 13 mitochondrial PCGs (length: 6.7 kbp) from 149 species sequenced in this study; and 2) the same mitochondrial genes from 282 species including mitogenomes downloaded from GenBank; 3) the 1164 nuclear markers (length: 1.13 Mbp) from 149 species sequenced in this study; and 4) the combined dataset of 13 mitochondrial and 1164 nuclear markers (length: 1.14 Mbp) from 285 species. We reasoned that the two mitochondrial datasets would allow us to assess how the increasing sampling improves the phylogeny and make a more direct comparison with published phylogenies that were mostly based on mitochondrial markers.

To evaluate whether our datasets have evolved under globally stationary, reversible, and homogeneous (SRH) conditions, we applied matched-pairs tests of homogeneity (Bowker’s test) implemented in SymTest v2.0.55 (https://github.com/ottmi/symtest) to test for potential compositional heterogeneity (Jermiin et al., 2004; Misof et al., 2014). Tests were applied on 1) the amino acid dataset, 2) the nucleotide dataset with all codon positions, and 3) the nucleotide dataset including first and second codon positions only. The results indicate that the amino acid dataset exhibited much less compositional heterogeneity compared to the nucleotide datasets, of which the dataset containing only the first and second codon positions showed less among-lineage heterogeneity than the one including all codon positions (Fig. S1). Additionally, the third codon position possibly suffers strong substitution saturation after >200 million years of evolution (Johnson et al., 2018). Taking together, we used the amino acid dataset and the nucleotide dataset including only the first and second codon positions for the subsequent phylogenetic tree inference.

All datasets were partitioned by genes. All phylogenies were inferred with the Maximum Likelihood (ML) approach in IQ-TREE2 v2.2.0.3 (Minh et al., 2020). For nucleotide datasets, all available DNA models were tested in IQ-TREE2. For amino acid datasets, selected protein models Q.insect, WAG, LG, and JTT (setting: -mset Q.insect, WAG, LG, JTT) were tested on nuclear datasets and the model mtInv (setting: -mset mtInv) was tested on mitochondrial datasets for computational efficiency. To decide on the partitioning scheme and the substitution model, we asked IQ-TREE2 to perform extended model selection on each gene with free rate heterogeneity followed by tree inference and subsequently merge two or more genes until the model fit does not increase any further (setting: -m MFP+MERGE; Chernomor et al., 2016; Kalyaanamoorthy et al., 2017). The best partitioning scheme and the best-fit models were chosen by the highest BIC (Bayesian Information Criterion) scores. Bootstrapping was conducted using the approximate likelihood ratio test (SH-aLRT) and ultrafast bootstrap methods with 1000 replicates as well as the approximate Bayes test (setting: -B 1000 --alrt 1000 --abayes; Anisimova et al., 2011; Anisimova and Gascuel, 2006; Hoang and Chernomor, 2017). All other setting options were kept default.

To account for the differences in the evolutionary history of genes, we inferred the species tree from gene trees using the Accurate Species Tree ALgorithm (ASTRAL-III) (Zhang, 2018). Specifically, we used Weighted ASTRAL (wASTRAL v1.15.2.3) to take into account phylogenetic uncertainty by integrating signals from branch length and branch support in gene trees (Zhang, 2022). We used the nucleotide alignment of each gene without the third codon position as the input to IQ-TREE2, which inferred each gene tree with 1000 replicates of ultrafast bootstrapping. Then, the combined tree file was analyzed in wASTRAL with 16 rounds of search and subsampling (setting: -R).

### Divergence time estimation with fossils

We used the nuclear dataset comprising the first and second codon positions for the molecular dating analyses. From all 1164 nuclear markers, we filtered out genes with less than 143 out of 149 (i.e. <95%) species present. This results in 57 markers. Then, we concatenated these alignments into one dataset. In addition, we made five gene subsets out of the 57 genes, each of which contains 11 to 12 randomly assigned markers. Each subset was concatenated into a separate dataset. We then decided on the best partitioning scheme and substitution model for each dataset in IQ-TREE2 (setting: -m MF+MERGE).

For node calibration, we assigned 30 fossils as calibration points throughout the tree (Table S2). The taxonomic placement of each fossil was determined based on the apomorphy of the corresponding taxon. Fossil taxa were selected based on their verified taxonomic and chronostratigraphic placement, to cover as low classification levels as possible, especially for the lineages present in molecular analyses. The fossil age was used as the minimum constraint of the node. Maximum constraints were chosen based on the phylogenetic placement, estimated dates from previously published studies, and the fossil history of certain clades. For the root, the oldest fossils assigned to the extant lineages of Fulgoromorpha are *Barremixius petrinus* and *Karebopodoides aptianus* of family Cixiidae, dated to the Lower Cretaceous, Barremian (125.77 - 121.4 Ma; Luo et al., 2021; Maksoud et al., 2017). Thus, we placed the minimum constraint of the root at the calibrated age of the two fossils (125 Ma). For the maximum constraint of the root, we referred to the extinct superfamilies Coleoscytoidea, Surijokocixioidea, and Fulgoridioidea, for which the oldest fossils are known from the Permian, Roadian (273.15 - 266.5 Ma), the Permian, Wordian (267.3 - 264.12 Ma), and the Lower Jurassic, Hettangian (201.6 - 199.2 Ma), respectively (Bucher et al., 2024; Szwedo, 2018; Szwedo et al., 2004). It is generally accepted that Coleoscytoidea is a side group, distantly related to the modern crown Fulgoromorpha, while Surijokocixioidea and Fulgoridioidea are considered as stem groups to the modern crown group (Bourgoin and Szwedo, 2023, 2022; Szwedo, 2018). However, whether Surijokocixioidea or Fulgoridioidea is more closely related to the crown group is still under debate (Bourgoin and Szwedo, 2023; Brysz and Szwedo, 2019; Szwedo, 2017). We chose the maximum constraint of the root at the base of Wordian (267.3 Ma) where the oldest fossil of Surijokocixioidea resides. Johnson et al. (2018) estimated the root age of crown Fulgoromorpha at ∼206 Ma with a 95% confidence interval (hereafter: CI) ranging from 178 to 241 Ma. Thus, our constraints can be considered relatively conservative (Table S2).

Regarding the maximum constraint for fossils in different modern families, the oldest fossils came from the Lower Cretaceous, Barremian (125.77 - 121.4 Ma) as mentioned above and all fossils from earlier periods, such as Jurassic (201.6 - 145 Ma), belong to the extinct, more ancient stem groups such as Fulgoridioidea. Thus, we placed the maximum constraint at the base of Jurassic (201.6 Ma) for fossils in families Cixiidae, Delphacidae, Kinnaridae, Achilidae, Derbidae, Fulgoridae, and Dictyopharidae. As for other relatively recently diverged families, since there are no clear patterns in fossil distribution, we referred to their host plants. Flowering plants, which are the main food source of all modern crown group planthoppers, first appeared in the fossil record in the early Cretaceous and began to diversify in the mid to later Cretaceous (Zuntini et al., 2024). It is very likely that the diversification of flowering plants drove the diversification of the derived planthopper families. Therefore, we placed the maximum constraint at the base of the Cretaceous (145 Ma) for families Tropiduchidae, Lophopidae, Nogodinidae, Issidae, Caliscelidae, and Ricaniidae (Table S2). The ages of all geologic periods were derived from the ICS International Chronostratigraphic Chart v2023/09 (http://www.stratigraphy.org/ICSchart/ChronostratChart2023-09.pdf).

We also applied a different set of maximum constraints based completely on previously inferred time trees. We referred to the work done by Johnson et al. (2018) where they produced a dated phylogeny for major Hemipteran clades including nine planthopper families. We chose the upper 95% confidence interval of the estimated age of the node that goes back one node deeper in the phylogeny as the maximum constraint for the target family. For example, the split between Acanaloniidae and Flatidae was dated to 76 Ma (95% CI: 58 - 96 Ma), and the split between the Acanaloniidae-Flatidae clade and Caliscelidae was dated to 90 Ma (95% CI: 70 - 109 Ma). Then, we chose 109 Ma as the maximum constraint for fossils in Flatidae and Acanaloniidae. Following this rule, we placed the maximum constraint at 241 Ma for fossils of Cixiidae, Delphacidae, Kinnaridae, Achilidae, Derbidae, Fulgoridae, and Dictyopharidae, 144 Ma for Tropiduchidae, and 123 Ma for Lophopidae, Nogodinidae, Issidae, Caliscelidae, and Ricaniidae. The maximum constraint of the root was set to 349 Ma as the upper 95% confidence interval of the estimated age of auchenorrhynchans (309 Ma, 95% CI: 275 - 349 Ma) (Table S2). The alternative constraints are more conservative for the root and relatively old families and less conservative for the derived families than our constraints based on fossil history and host plants.

The molecular dating analyses were performed in BEAST v2.7.6 (Bouckaert and Vaughan, 2019). We placed monophyly constraints on the nodes where both SH-aLRT and ultrafast bootstrap support were >95. For each dataset, the partition and its site model were set according to the best scheme found by IQ-TREE2 and were estimated later in BEAST. The rates of the branches were modeled under the optimized relaxed clock (Douglas et al., 2021). The uniform prior was applied to the root. For calibration points, we applied three different priors: uniform prior and log-normal prior with two different standard deviations (S = 1.0 or 0.5). For log-normal prior, we gave an offset equal to the minimum age and an adjusted mean (M) placing the maximum age at the 97.5% percentile. Other priors were kept default. Each dataset was run three times independently on each prior setting with a Markov Chain Monte Carlo (MCMC) chain length of at least 50 million to ensure good mixing. In addition, a run without sequence data (‘sample from prior’ in BEAST) was also performed for each dataset to examine the effective priors, compared to the specified priors. MCMC results were accessed with Tracer v1.7.2 (Rambaut et al., 2018). The verified log and tree files of the three independent runs using the same dataset were combined using LogCombiner v2.7.4 with 10% burnin. The maximum clade credibility tree of each dataset with mean node heights was summarised from the combined tree set by TreeAnnotator v2.7.4. All trees were visualized in FigTree v1.4.4 (http://tree.bio.ed.ac.uk/software/figtree/). The node ages were extracted using the *treeio* R package (Wang et al., 2019). The summary of node ages was illustrated using Processing 3 (http://www.processing.org). All figures were edited in Inkscape (https://inkscape.org).

## Results

### Metagenomic data summary and phylogeny overview

From 149 metagenomes, we obtained between 5.3 and 34.6 Gb of data (16.3 Gb on average) corresponding to 35.3 and 230.4 million reads (Table S3). The metagenomes collectively enabled the assembly of 1164 markers out of 2510 single-copy nuclear genes from the BUSCO hemipteran database. Among these 1164 markers, we obtained 91 to 1161 markers (1026 on average) from each metagenome. One sample was represented by <10% of markers and five samples were represented by <20% of markers The phylogeny based on 1164 nuclear markers provided the most robust family tree of planthoppers so far, showing full support on most branches (Fig. S4). The addition of species with mitogenomes extracted from GenBank did not change the topology of our best tree while contributing more details to the relationships within each family (Figs. 1 & S5). However, with this expanded dataset, we also saw more weakly-supported shallow branches and encountered the failure of convergence of non-parametric ultrafast bootstrap (Fig. 1; Supplementary material Data file S5). One possible reason is that the additional mitogenomes, with a number almost equal to the metagenomes on the phylogeny, brought too little additional phylogenetic information while the algorithm tried to solve a more complex tree space. This made species with only mitogenomes prone to phylogenetic artifacts (e.g., long-branch attraction) after diverging from each other hundreds of million years ago. Indeed, compared to the best tree, the trees based on solely 13 mitochondrial PCGs recovered different, partially supported placement of a few families (Figs. S2-3). The topologies were more similar to the best tree when more samples were added, although the changed topologies were weakly supported and several families were still misplaced (Figs. S2-3; Wang et al., 2023).

**Figure 1.**
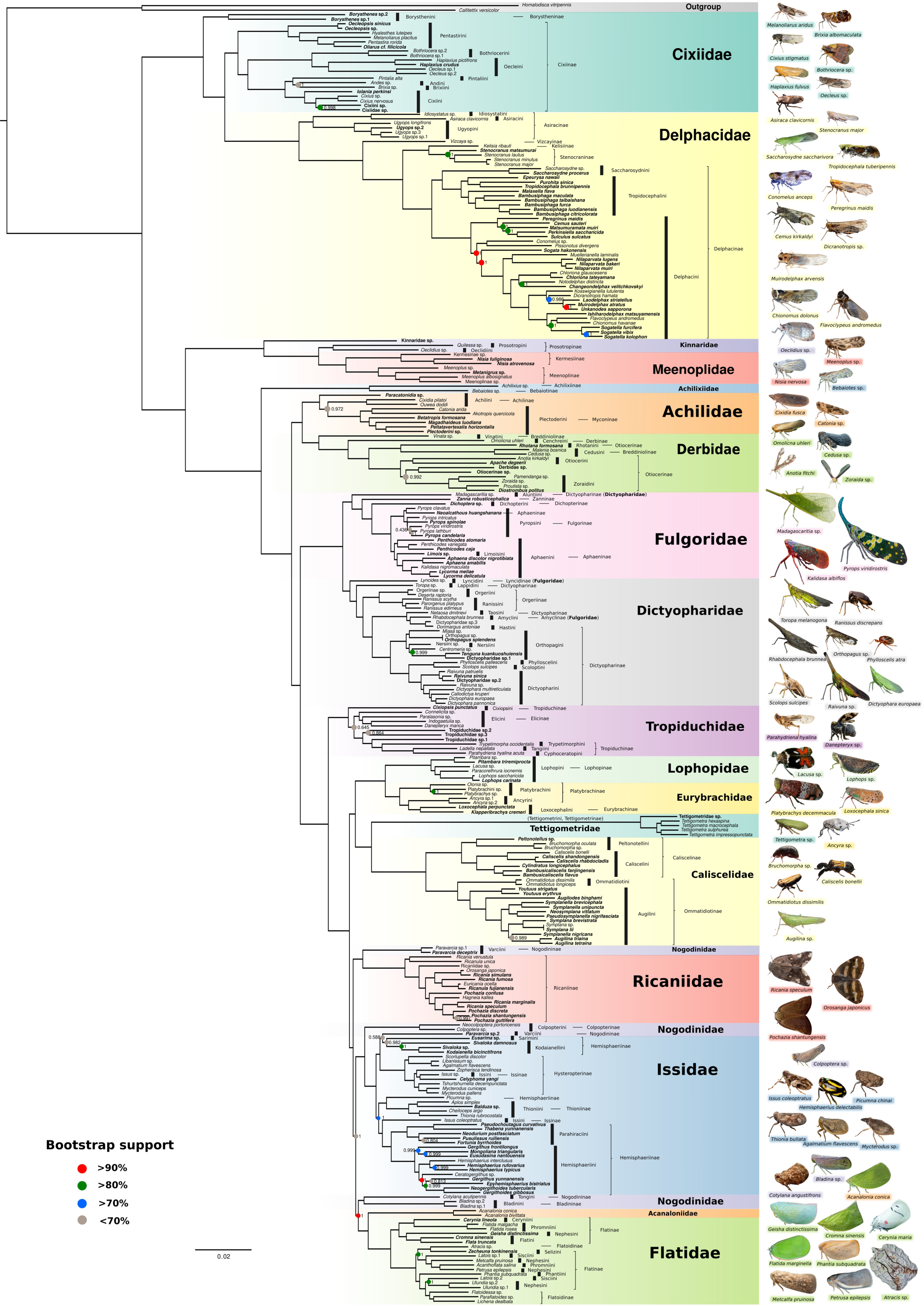
The best Maximum-Likelihood tree based on the concatenated nucleotide dataset of 1164 nuclear and 13 mitochondrial markers of 285 planthopper species. The SH-aLRT branch support values <95 are represented by colored circles. Higher or full support values are not shown. Ultrafast bootstrap supports are not shown due to the failure of convergence. The support from the approximate Bayes test is shown next to each node. Species contributing only mitogenomes have their names in bold. The tribe and subfamily information are indicated next to each species according to Fulgoromorpha Lists On the Web (FLOW; Bourgoin, 2024). All photos of planthoppers are taken from iNaturalist.org, after verification that licenses permit sharing and adaptation; species and observers’ IDs are listed in Table S4.

The best-obtained topology was largely similar when analyzing protein or DNA alignments. Both approaches yielded strongly supported, highly similar topologies, except for a few rogue taxa with uncertain placement in all trees (Supplementary material Data files S3-5). When only mitochondrial markers were used, both approaches resulted in weakly supported topologies that differed significantly from each other, indicating that mitogenomic data are insufficient for resolving deep phylogenetic relationships in our case (Supplementary material Data files S1-2). Furthermore, the species tree inferred from nuclear gene trees using ASTRAL mostly agrees with the best tree, although a few key differences exist (Fig. S6).

In the following sections, we present novel findings of planthopper phylogeny based on our best trees. The above-mentioned differences between topologies inferred from different datasets or models are described in detail.

### The superfamily Delphacoidea

All our inferred trees strongly supported the recently recognized superfamily Delphacoidea, which includes the families Cixiidae and Delphacidae (Figs. 1 & S2-5; Bourgoin and Szwedo, 2023). Delphacoidea is sister to the rest of the crown Fulgoromorpha. Within the family Cixiidae, our analyses strongly supported the monophyly of the subfamily Cixiinae. However, the two species from the genus *Borysthenes* (subfamily Borystheninae) were placed outside the monophyletic group formed by the remaining Cixiids – all from Cixiinae – and the family Delphacidae (Figs. 1 & S3). The origin of the *Borysthenes* clade predated the separation between Cixiinae and Delphacidae, arguing for its taxonomic classification as a new family separate from Cixiidae (see “Taxonomic decisions” section).

The monophyly of Delphacidae was also recovered (Figs. 1 & S2-5). The subfamily Asiracinae, including species from genera *Ugyops*, *Asiraca*, and *Idiosystatus*, is sister to the rest of Delphacidae. Following Asiracinae is the clade comprising the subfamilies Vizcayinae, Kelisiinae, and Stenocraninae, with the latter two sister to each other and the monophyletic subfamily Delphacinae sister to this clade. The three tribes within Delphacinae – Saccharosydnini, Tropidocephalini, and Delphacini – were also recovered as monophyletic and well-supported.

### Kinnaridae and Meenoplidae

All analyses recovered the monophyletic clade of the families Kinnaridae and Meenoplidae, which is sister to the rest of Fulgoroidea (Figs. 1 & S2-5). Their age and phylogenetic distance from the remaining fulgoroid families argued for its classification as a new superfamily (see “Taxonomic decisions” section). Regarding Meenoplidae, our inferred trees strongly supported its monophyly with an inclusion of four species from the subfamily Meenoplinae. For Kinnaridae, the monophyly was recovered for the two species *Quilessa* sp. and *Oeclidius* sp. representing the subfamily Prosotropinae. However, the unrecognized Kinnaridae sp. from Wang et al. (2023) was placed outside the monophyletic group formed by the other kinnarids and meenoplids, suggestive of misidentification or the paraphyly of Kinnaridae (Figs. 1 & S3).

### Achilixiidae, Achilidae, and Derbidae

We strongly supported the monophyly of Achilixiidae by including species from the two known genera *Achilixius* and *Bebaiotes* (Figs. 1 & S2-5). Additionally, we placed Achilixiidae outside of the Achilidae-Derbidae clade, as sister to the rest of the superfamily Fulgoroidea excluding Kinnaridae and Meenoplidae. Achilidae and Derbidae were reciprocally monophyletic, i.e. they were sister groups (Figs. 1 & S2-5). Note that Wang et al. (2023) placed the sample identified as meenoplid *Suva longipenna* within Derbidae. By sampling more broadly, we firmly placed this species – renamed as Otiocerinae sp. – within the derbid subfamily Otiocerinae, suggesting that ‘*Suva’ longipenna* was misidentified and wrongly placed (see “Taxonomic decisions” section). Furthermore, our inferred trees placed Achilidae and Derbidae as sister to the remaining fulgoroids, with strong support (Figs. 1 & S4-5). This topology was different from the ones based on only mitogenomes, which switched positions of the Achilidae-Derbidae clade and the Fulgoridae-Dictyopharidae clade on the nuclear trees and recovered Fulgoridae and Dictyopharidae as sister to the remaining families including Achilidae and Derbidae (Figs. S2-3). However, the support for this alternative topology was weak.

### Fulgoridae and Dictyopharidae

Our inferred trees clearly recovered two monophyletic groups, representing Fulgoridae and Dictyopharidae, and their sister relationship (Figs. 1 & S4-5). Interestingly, *Lyncides* sp. and *Madagascaritia* sp., which are only found on Madagascar and neighboring islands, were both sister to the rest of the sampled lineages in their respective families (*Lyncides* sp. in Dictyopharidae and *Madagascaritia* sp. in Fulgoridae). These placements suggest that Gondwanaland, of which Madagascar was part during the pre-Cretaceous times, might be the birthplace of the ancestors of all modern fulgorids and dictyopharids.

Additionally, our phylogenies suggested misplacements of several taxa. For example, the above-mentioned *Lyncides* sp. and *Madagascaritia* sp. were placed within Dictyopharidae and Fulgoridae, respectively. This is in contrast with the current taxonomic placement of *Lyncides* sp. as a fulgorid and *Madagascaritia* sp. as a dictyopharid. Similarly, *Rhabdocephala brunnea*, currently classified as a fulgorid, was strongly placed within Dictyopharidae. The genus *Zanna*, which was represented by one species *Z. robusticephalica* in our study and was recovered more closely allied with Dictyopharidae in Urban and Cryan (2009), was firmly placed inside Fulgoridae.

### Tropiduchidae, Tettigometridae, Caliscelidae, Lophopidae, and Eurybrachidae

We recovered the monophyly of Tropiduchidae and its sister relationship to the remaining families of Fulgoroidea (Figs. 1 & S2-5). The placement of *Cixiopsis punctatus* (tribe Cixiopsini) was unresolved based solely on its mitogenome (Figs. 1 & S3, 5).

Tettigometridae was recovered as a sister family to Caliscelidae (Figs. 1 & S4-5). The sister relationship between Lophopidae and Eurybrachidae was also strongly supported. In addition, we found a sister relationship between the Tettigometridae-Caliscelidae clade and the Lophopidae-Eurybrachidae clade, recovering a broader monophyletic group consisting of these four families (Figs. 1 & S4-5). Our inferred trees based on mitochondrial PCGs recovered a similar topology but failed to place Tettigometridae within the monophyletic clade of Lophopidae, Eurybrachidae, and Caliscelidae (Figs. S2-3). However, the placements of Tettigometridae were always close to the other three families. Interestingly, the ASTRAL species tree inferred from thousands of nuclear gene trees broke up the four-family monophyly, placing the Tettigometridae-Caliscelidae clade as sister to the rest, and then the Lophopidae-Eurybrachidae clade as sister to the remaining clades (Fig. S6).

Tettigometridae showed a very long branch, suggesting a faster rate of molecular evolution, which might be linked to its specialized symbiotic relationship with ants (Dejean et al., 2000; Lehouck et al., 2004). Within Caliscelidae, the monophyly of two subfamilies (Caliscelinae and Ommatidiotinae) and the four tribes (Peltonotellini, Caliscelini, Ommatidiotini, and Augilini) were well-supported.

### The partially resolved relationships among the rest of the families

Despite using >1000 nuclear markers, we were not able to fully resolve the relationships among the remaining families, including Nogodinidae, Ricaniidae, Issidae, Acanaloniidae, and Flatidae. However, our inferred trees still provided insights into some key relationships among them.

For the family Nogodinidae, we confirmed its paraphyly and suggested its strong links to other families. The species from the genus *Paravarcia* (tribe Varciini, subfamily Nogodininae) was strongly placed sister to Ricaniidae, whose monophyly was well-recovered (Figs. 1 & S4-5). Similarly, the two species *Neocolpoptera portoricensis* and *Colpoptera* sp. from the tribe Colpopterini (subfamily Colpopterinae) were sister to Issidae, whose monophyly was also fully supported. Additionally, *Cotylana acutipennis* (tribe Tongini, subfamily Nogodininae) was sister to the other two Nogodinidae species from the genus *Bladina* (tribe Bladinini, subfamily Bladininae). Among these sister groups, however, the relationships were uncertain. Note that *Paravarcia* sp.2 – sister to *Paravarcia deceptrix* in Wang et al. (2023) – was not grouped with the other *Paravarcia* species and its position is uncertain (Figs. 1 & S3).

The sister relationship between Acanaloniidae and Flatidae was recovered with strong support (Figs. 1 & S4-5). Note that Wang et al. (2023) placed Acanaloniidae, represented by a single species *Acanalonia* sp. sampled from China, sister to Tropiduchidae. This is highly skeptical as there are no records of Acanaloniidae in China or East Asia, and thus the specimen seems to have been misidentified. Our analyses firmly placed this species – renamed as Tropiduchidae sp.3 – within the family Tropiduchidae.

Within Issidae, the monophyly of the subfamilies Hysteropterinae and Thioniinae was well-supported (Figs. 1 & S4-5). Note that the specimen identified as *Issus* sp. (subfamily Issinae) sampled from Kazakhstan was placed within the subfamily Hysteropterinae, which might be due to specimen misidentification. The subfamily Hemisphaeriinae was recovered as paraphyletic, although the support was weak. Within Hemisphaeriinae, the monophyly of tribes Hemisphaeriini and Parahiraciini and their sister relationship were well-supported. Furthermore, the species from the genus *Picumna*, currently classified within Hemisphaeriinae, was strongly supported as a sister group to Thioniinae, with species sampled exclusively from the tribe Thioniini. Within Flatidae, the subfamilies Flatinae and Flatoidinae, as well as the tribes proposed, were largely paraphyletic, indicating that the taxonomy within Flatidae requires a comprehensive investigation and revision in the future.

### The origin of the extant Fulgoromorpha was dated to at least the middle Permian

The first set of analyses applied uniform fossil priors with maximum age constraints based on fossil history and host plants. This resulted in a root age of 263 Ma (95% CI: 255 - 267 Ma) for the crown Fulgoromorpha, dating back to Guadalupian, Permian (Figs. 2 & S7).

**Figure 2.**
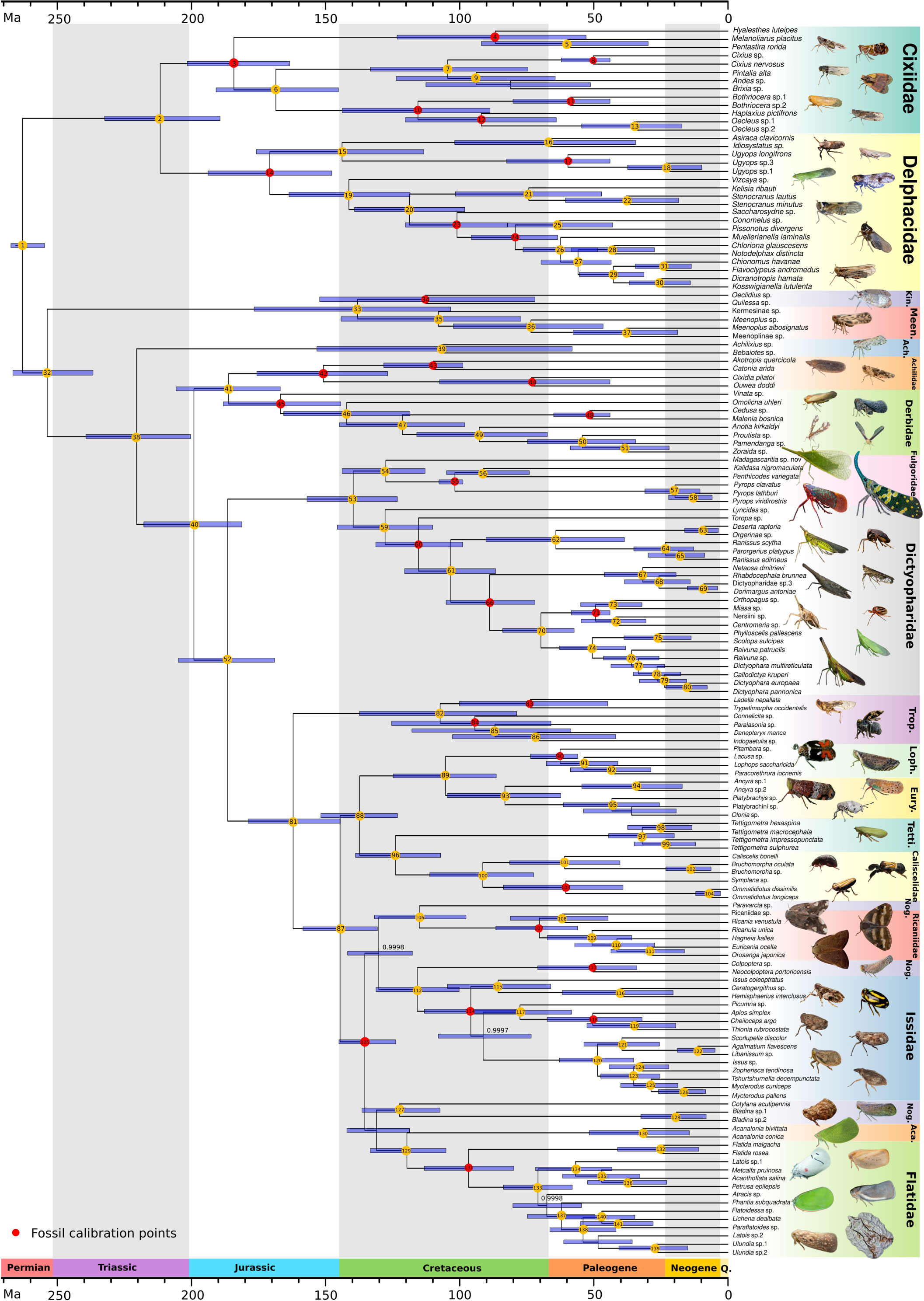
The dated phylogeny of 149 planthopper species estimated in BEAST, based on a reduced dataset of 57 nuclear genes and the topology of the best Maximum-Likelihood tree from IQ-TREE2 in Figure S4. All 30 calibration points with fossils are indicated by red circles and were assigned with uniform priors and our age constraints based on fossil history and host plants. The nodes fixed in BEAST analyses are labeled inside colored circles. The unfixed nodes are not labeled. The 95% confidence intervals of node age are noted as light blue bars. Bayesian posterior probabilities that are less than one are shown next to the nodes. Timescales are in millions of years. Family names are abbreviated as follows: Kinnaridae (Kin.), Meenoplidae (Meen.), Achilixiidae (Ach.), Tropiduchidae (Trop.), Lophopidae (Loph.), Eurybrachidae (Eury.), Tettigometridae (Tetti.), Nogodinidae (Nog.), and Acanaloniidae (Aca.).

Within the crown Fulgoromorpha, we estimated the divergence between Cixiidae and Delphacidae, or the age of the extant superfamily Delphacoidea, at 212 Ma (95% CI: 189 - 232 Ma) – during the Upper Triassic. The extant lineages of Cixiidae and Delphacidae started diversifying at 184 Ma (95% CI: 163 - 202 Ma) and 171 Ma (95% CI: 148 - 194 Ma), respectively. The age of the superfamily Fulgoroidea, including Kinnaridae and Meenoplidae, was estimated at 254 Ma (95% CI: 237 - 267 Ma) – the Late Permian/Early Triassic, which also marks the divergence of Kinnaridae and Meenoplidae from other Fulgoroidea families. Next, Achilixiidae diverged from the remaining Fulgoroidea families at 220 Ma (95% CI: 200 - 239 Ma). Achilidae and Derbidae diverged from the rest of the Fulgoroidea families at 199 Ma (95% CI: 181 - 218 Ma) and diverged from each other at 186 Ma (95% CI: 167 - 206 Ma). Then, Fulgoridae and Dictyopharidae diverged from the remaining families at 187 Ma (95% CI: 169 - 205 Ma) and diverged from each other at 140 Ma (95% CI: 123 - 157 Ma). The age of crown Fulgoridae and Dictyopharidae was similar, dating back to ∼128 Ma. For the remaining families, Tropiduchidae diverged from the other families at 162 Ma (95% CI: 144 - 179 Ma). Then, the monophyletic group consisting of Lophopidae, Eurybrachidae, Tettigometridae, and Caliscelidae diverged from the rest of the families at 145 Ma (95% CI: 131 - 158 Ma). Within this group, Lophopidae and Eurybrachidae diverged from the sister group of Tettigometridae and Caliscelidae at 137 Ma (95% CI: 123 - 152 Ma) and diverged from each other at 105 Ma (95% CI: 86 - 125 Ma). Tettigometridae diverged from Caliscelidae at 124 Ma (95% CI: 107 - 139 Ma). The root age of the remaining families, including Ricaniidae, Nogodinidae, Issidae, Acanaloniidae, and Flatidae, was dated to 135 Ma (95% CI: 124 - 145 Ma). As the relationships among these five families are uncertain, we only provide estimates for the crown age of some families. The extant lineages of Ricaniidae and Issidae started diversifying at 70 Ma (95% CI: 56 - 87 Ma) and 96 Ma (95% CI: 78 - 113 Ma), respectively. Acanaloniidae diversified from Flatidae at 120 Ma (95% CI: 105 - 133 Ma).

### Alternative fossil priors and age constraints yielded different node ages

We first compared the posterior and marginal prior distributions of calibrated nodes resulting from different fossil priors – uniform, log-normal (S=1), or log-normal (S=0.5) – under the same constraints based on fossil history and host plants (Figs. 3-4). Note that the marginal prior distributions, which resulted from the interactions between all calibration priors and tree prior, were not the same as the defined fossil priors. Across our dated phylogeny, many calibrated nodes showed shifts in posterior distributions from marginal priors, indicating that molecular data provided additional information about their age. Among these nodes, the uniform and the flat log-normal (S=0.5) priors often yielded the oldest estimates, while the steep log-normal (S=1) prior yielded the youngest (Figs. 3-4; Supporting Information Data S7). Other nodes (nodes 11, 17, 44, 83, 107, and 113) showed largely overlapped posterior and prior distributions, suggesting that our molecular data lacked sufficient information to guide the posterior away from the prior and give meaningful predictions on node ages. Regarding the root of Fulgoromorpha, we applied the uniform prior to the entire analysis, no matter what fossil priors were used. All three runs resulted in similar ages approaching the maximum constraint, indicating that the root constraint might still be strict and the origin of crown Fulgoromorpha is possibly older than 267.3 Ma (Fig. 3), as suggested by fossil records and current classification proposals (Bourgoin and Szwedo, 2023, 2022; Bucher et al., 2024).

**Figure 3.**
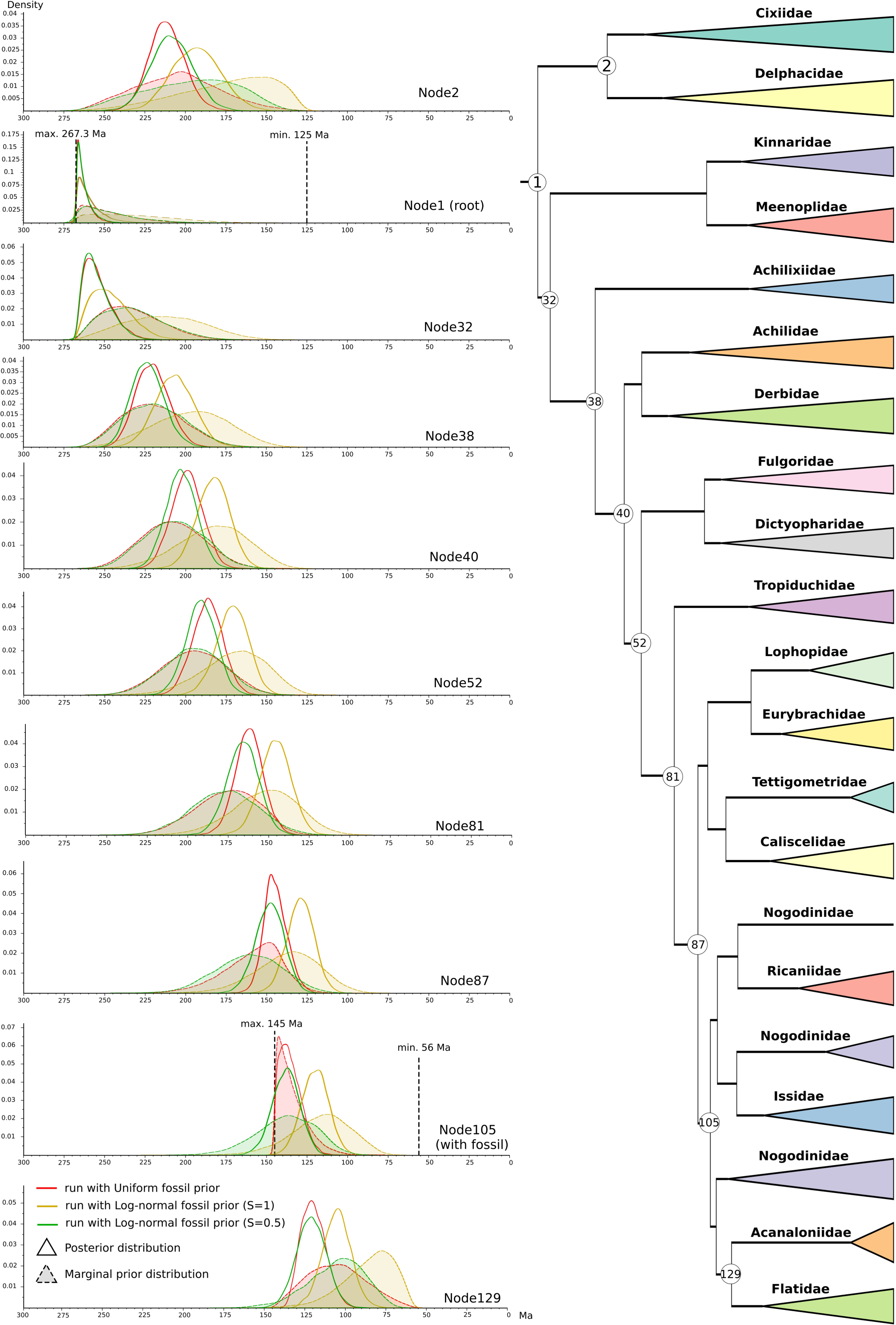
Marginal prior (shaded area under the dotted curve) and posterior (unshaded area under the solid curve) distribution of the age estimates of ten selected nodes from analyses with uniform (red), log-normal (S=1, yellow), or log-normal (S=0.5, green) fossil priors. The minimum and maximum age constraints on the root (node1) and one fossil (node105) are noted as vertical dotted lines. We applied uniform prior to the root through entire runs, no matter what fossil priors were used. The collapsed topology from Figure 2 is shown on the right.

**Figure 4.**
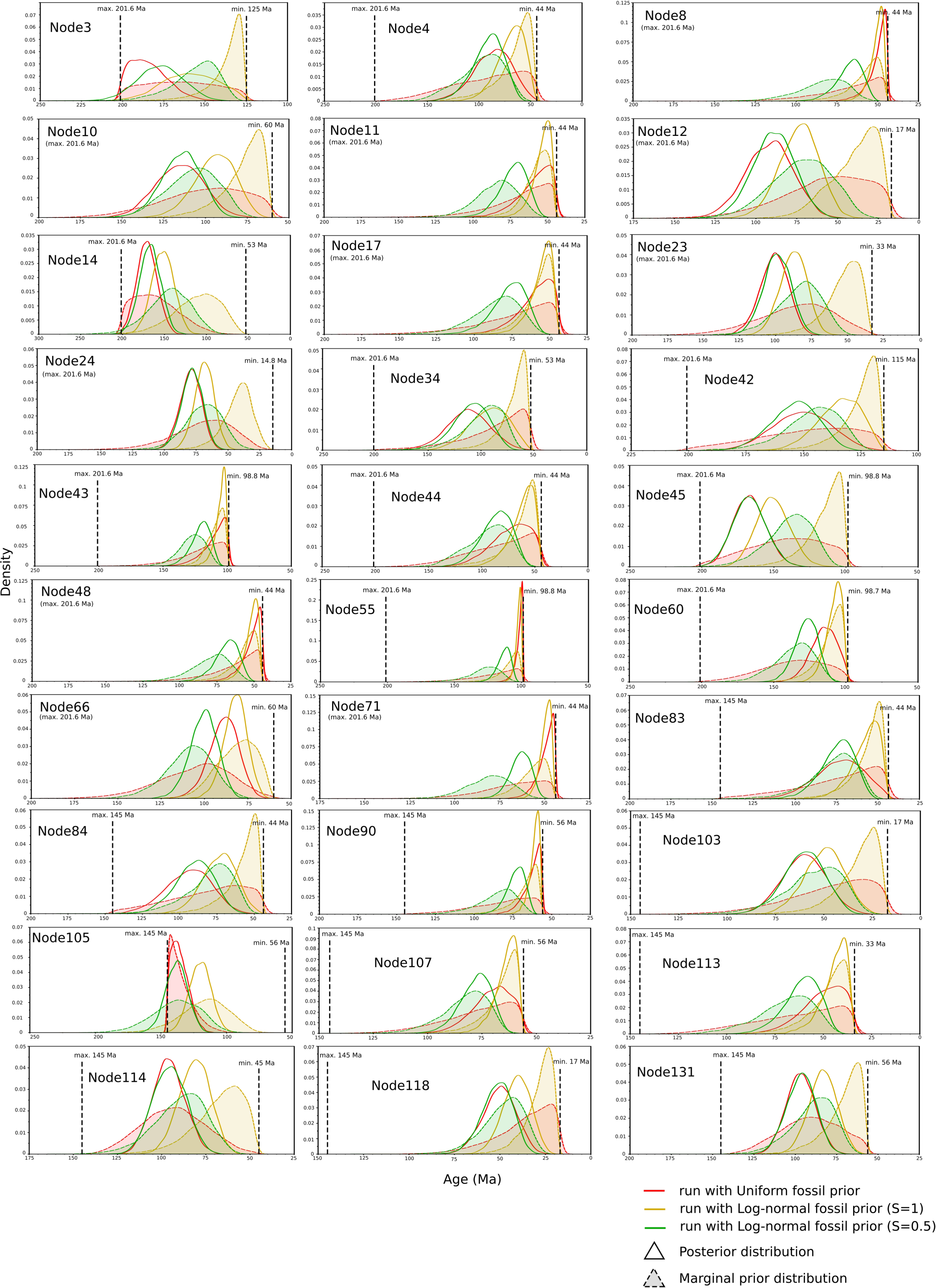
Marginal prior (shaded area under the dotted curve) and posterior (unshaded area under the solid curve) distribution of the age estimates of all 30 fossils from analyses with uniform (red), log-normal (S=1, yellow), or log-normal (S=0.5, green) fossil priors. The minimum and maximum age constraints on each fossil are noted as vertical dotted lines. The out-of-range maximum ages are shown below node labels.

We then compared the node ages resulting from two different sets of constraints under the same uniform prior, one based on fossil history and host plants and the other based on previously published dated phylogeny (Fig. 5). Interestingly, the node ages were mostly consistent under different constraints, despite the alternative constraints being more conservative for the relatively old lineages and less for the recently diversified lineages. The nodes showing significant age differences were all deep nodes sitting away from the tips. The posterior and marginal prior distributions showed that their ages shifted in the same direction as the marginal priors (Fig. 6). When an older maximum age was applied, we observed shifts of both the marginal prior and posterior distribution towards older ages with a wider CI. The opposite trend was observed when a younger maximum age was used. The largest shifts were observed in node1 and node32, which represented the root of crown Fulgoromorpha and Fulgoroidea, respectively.

**Figure 5.**
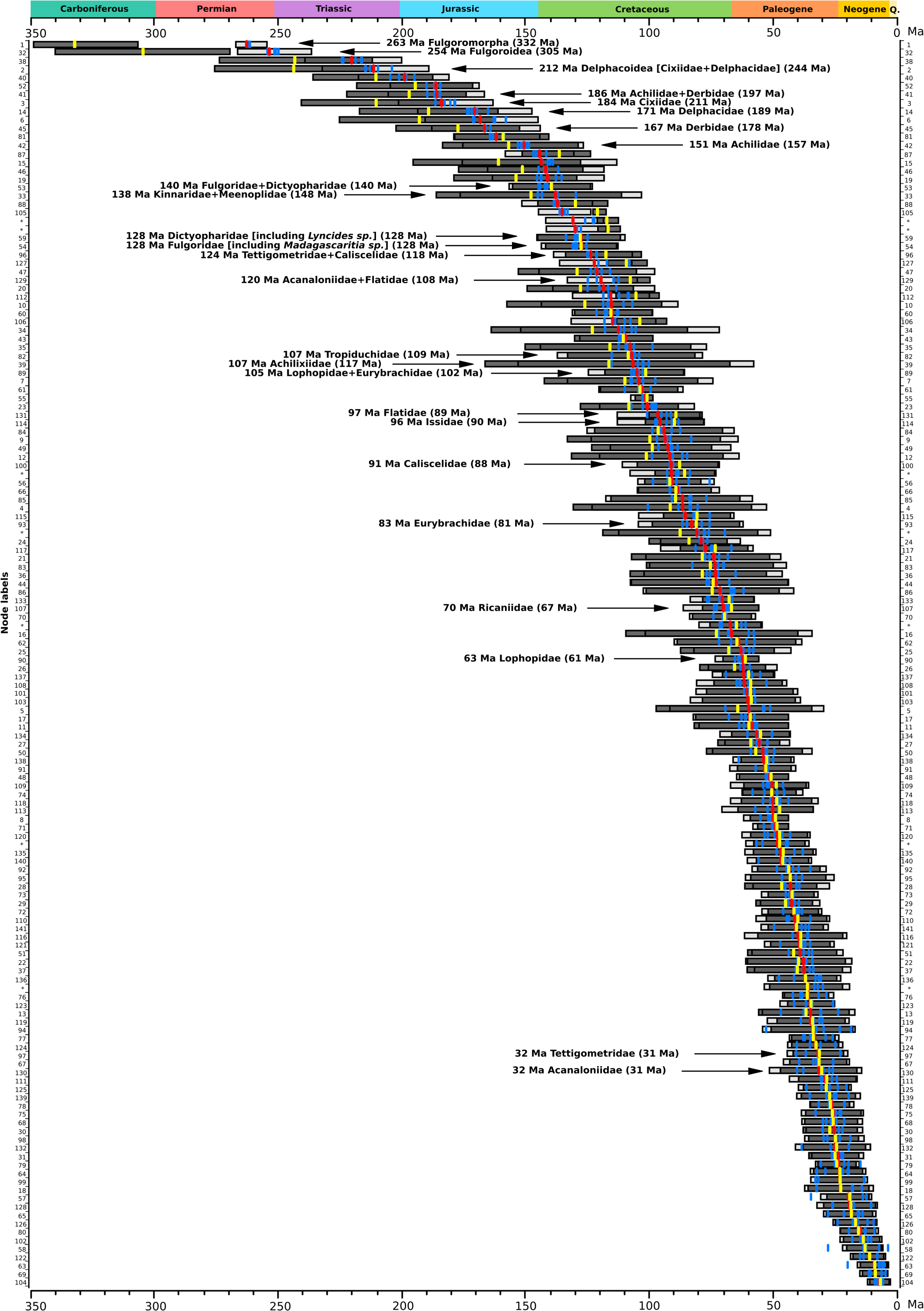
Sorted node age estimates from analyses under uniform fossil priors and different sets of maximum age constraints. The light and dark grey horizontal bars represent the 95% confidence interval of age estimates under our constraints based on fossil history and host plants (Max. root age: 267.3 Ma) and alternative constraints based on the published dated tree from Johnson et al. (2018) (Max. root age: 349 Ma), respectively. Mean ages are noted as red and yellow bars in the middle, respectively. Additionally, we show the mean ages (blue bars) derived from independent analyses of five sub-groups of genes. These analyses were only performed under our age constraints. The mean ages of selected families and clades are shown next to the bar, with ages estimated under alternative constraints in the brackets. Node labels corresponding to the ones in Figure 2 are shown on the left and right sides of the figure. Unfixed nodes are indicated with asterisks.

**Figure 6.**
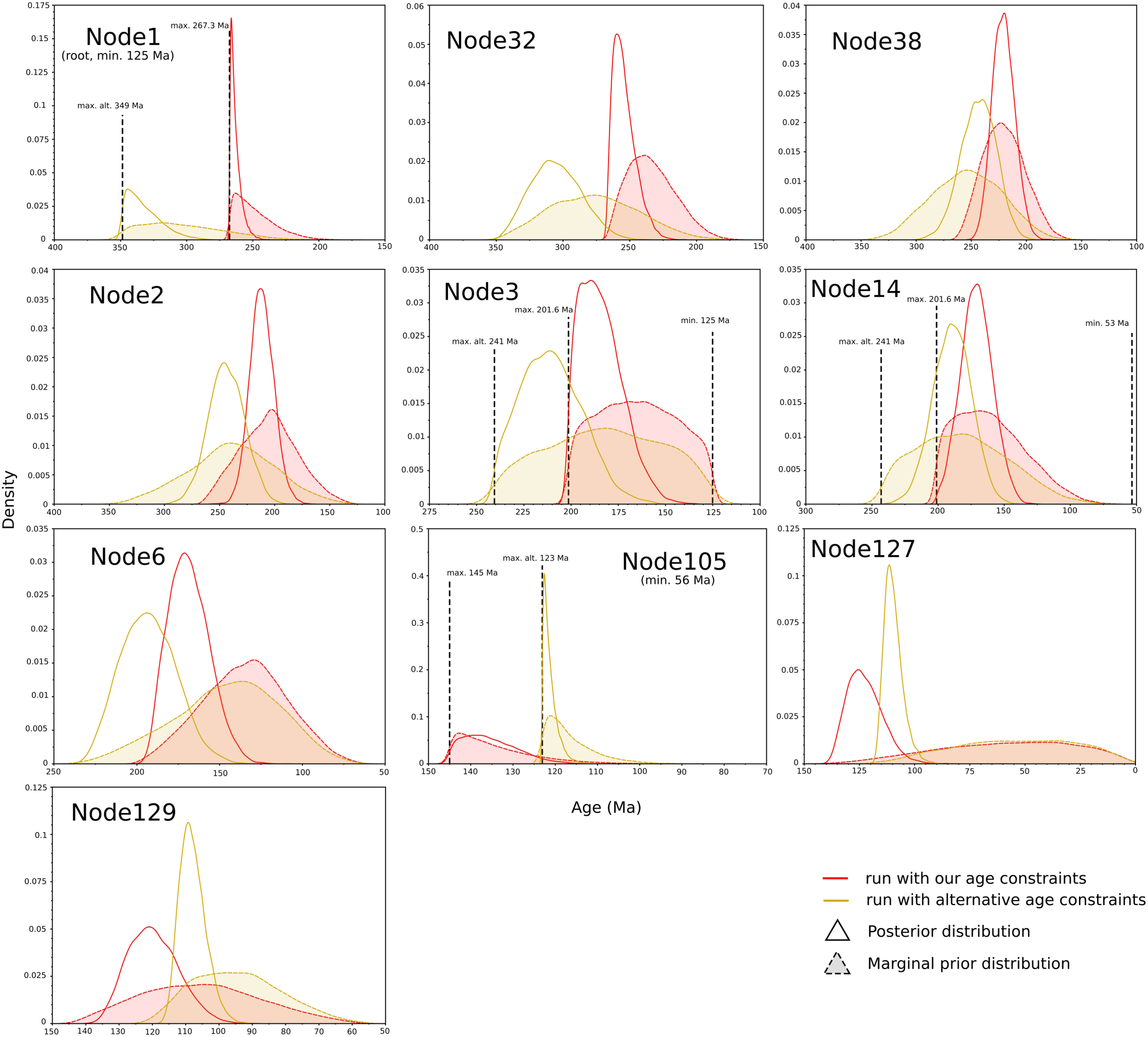
Marginal prior (shaded area under the dotted curve) and posterior (unshaded area under the solid curve) distribution of the age of nodes showing significant shifts under different sets of age constraints. The minimum and maximum age constraints are noted as vertical dotted lines. The alternative maximum constraints are indicated as “max. alt.”. The out-of-range maximum ages are shown below node labels.

Their posterior distributions almost showed no overlaps to the ones under a younger maximum age. Although the distributions are flatter, the posterior clearly shifted away from the marginal prior towards older age, indicating that the real ages of Fulgoromorpha and Fulgoroidea might be older than what we observed under our age constraints. The ages estimated under the alternative constraints placed the origin of crown Fulgoromorpha (332 Ma, 95% CI: 307 - 349 Ma) in Mississippian, Carboniferous, and Fulgoroidea – including Kinnaridae and Meenoplidae – (305 Ma, 95% CI: 270 - 340 Ma) close to Carboniferous-Permian boundary.

## Taxonomic decisions

### 1. A new superfamily Meenoploidea Fieber, 1872 superfam. nov

Based on molecular branching and clear separateness of clade containing representatives placed in families Kinnaridae Muir, 1925 and Meenoplidae Fieber, 1972, from the other fulgoroid families, the new superfamily Meenoploidea Fieber, 1872 **superfam. nov.** is here proposed.

### Superfamily Meenoploidea Fieber, 1872 superfam. nov

#### Diagnostic characters

Median ocellus generally present; Presence of special wax gland pores on abdominal tergites 6-8 in female adults. Body strongly compressed; the tegmina holds tectiform. Pronotum short, broader than the head; mesonotum broader than the pronotum. Male genitalia with the genital plates large and complex; female genitalia with ovipositor modified, shortened, or reduced.

#### Content

**Family Kinnaridae Muir, 1925 stat. rev.**, Meenoplidae Fieber, 1872, **stat. rev.**

Family Kinnaridae Muir, 1925

#### Diagnostic characters

Costal cell wide; Costal vein of fore wing weakened in distal part like peripheral vein of wing membrane; nodal break present; Branches RP and MP with anastomosis at nodal level; Common claval vein reaching CuP before claval apex (achilidoid clavus); Frons strongly expanded laterad towards frontoclypeal suture; genal suture with a subantennal cross ridge on which a row of spinous processes is located and a laterally directed, conical process between the subantennal cross ridge and antenna; male pygofer with a slender, elongate, dorsoposteriorly directed process, ‘the pygofer appendage.

#### Content

The family Kinnaridae is limited to comprise only subfamily Kinnarinae Muir, 1925 with tribe Kinnarini Muir, 1925 and a single genus *Kinnara* Distant, 1906 (type species *Plemora ceylonica* Melichar, 1903), with 29 species (*K. albiabdominalis* Wang, Chou et Yuan, 2001; *K. albipennis* Liang, 2002; *K. bakeri* Muir, 1922; *K. brunnea* Muir, 1913; *K. ceylonica* (Melichar, 1903); *K. doto* Fennah, 1978; *K. flavifrons* Muir, 1913; *K. flavimarginalis* Wang, Chou & Yuan, 2001; *K. flavipes* (Bierman, 1907); *K. flavofasciata* Distant, 1916; *K. fulva* Muir, 1913; *K. fumata* (Melichar, 1903); *K. fusca* Liang, 2002; *K. guilinensis* Wang, Chou & Yuan, 2001; *K. maculata* Distant, 1912; *K. marginalis* Muir, 1922; *K. mediprocessa* Wang, Chou & Yuan, 2001; *K. nigrocacuminis* Muir, 1923; *K. nigrolineata* Muir, 1922; *K. ochracea* Liang, 2002; *K. penangensis* Muir, 1922; *K. sabahensis* Liang, 2002; *K. sordida* Muir, 1913; *K. spectra* Distant, 1912; *K. tricarinata* Wang, Chou et Yuan, 2001; *K. trifurcata* Wang, Chou et Yuan, 2001; *K. trimaculata* Liang, 2002; *K. unimaculata* Liang, 2002; and *K. vientianensis* Liang, 2002)

### Family Meenoplidae Fieber, 1872, stat. rev

#### Diagnostic characters

One or both claval veins typically granulated; RP and MP connected with crossvein rp-mp; Head with the vertex and frons (usually) broad and the lateral carinae strongly elevated; antennae generally short and simple - sensory plate organs star-like; Last segment of the labium elongate; Tegulae large.

#### Content

Prosotropinae Fennah, 1945, including genus *incertae sedis Kinnapotiguara* Xing, Hoch & Chen, 2013 (likely belong to Prosotropini), and tribes: Kinnocciini Emeljanov, 2006 (genera: *Iuiuia* Hoch & Ferreira, 2016; *Kinnacana* Remane, 1985; *Kinnoccia* Remane, 1985); Oeclidiini Emeljanov, 2006 (genera: *Micrixia* Fowler, 1904; *Oeclidius* Van Duzee, 1914; *Southia* Kirkaldy, 1904), Adolendini Emeljanov, 1984 (genera: *Adolenda* Distant, 1911; *Bashgultala* Dlabola, 1957; *Nesomicrixia* Emeljanov, 1984; *Paramicrixia* Distant, 1911; *Perloma* Emeljanov, 1984; *Valenciolenda* Hoch & Sendra, 2021); and Prosotropini Fennah, 1945 (genera: *Apocathema* Emeljanov, 2017; *Atopocixius* Muir, 1926; *Dineparmene* Fennah, 1945; *Eparmenoides* Fennah, 1945; *Lomagenes* Fennah, 1945; *Mauriciana* Campodonico, 2018; *Microissus* Fennah, 1947; *Oreopenes* Ramos, 1957; *Prosotropis* Uhler, 1895; *Quilessa* Fennah, 1942); Kermesiinae Kirkaldy, 1906 (genera: *Anorhinosia* Bourgoin, 1997; *Caledonisia* Bourgoin, 1997; *Eponisia* Matsumura, 1914; *Eponisiella* Emeljanov, 1984; *Fennahsia* Bourgoin, 1997; *Glyptodonisia* Bourgoin, 1997; *Insulisia* Bonfils & Attié, 1998; *Kermesia* Melichar, 1903; *Koghisia* Bourgoin, 1997; *Muirsinia* Bourgoin, 1997; *Nisamia* Emeljanov, 1984; *Nisia* Melichar, 1903; *Phaconeura* Kirkaldy, 1906; *Robigalia* Distant, 1916; *Suva* Kirkaldy, 1906; *Suvanisia* Bourgoin, 1997; *Tyweponisia* Bourgoin, 1997); and Meenoplinae Fieber, 1872 (genera: *Afronisia* Wilson, 1988; *Anigrus* Stål, 1866; *Distantiana* Bourgoin, 1997; *Meenoplus* Fieber, 1866; *Metanigrus* Tsaur & Yang, 1986; *Tsingya* Hoch & Bourgoin, 2014).

### 2. An elevation of the rank of Borysthenini Emeljanov, 1989 to family rank – Borysthenidae Emeljanov, 1989 stat. rev

Borysteninae as a subfamily of Cixiidae was proposed by (Emeljanov, 1989), based on the morphological peculiarities of its sole genus *Borysthenes* Stål, 1866 (Emeljanov, 1989; Liang, 2005). More recently, Luo et al. (2021) downgraded its rank to tribe in Cixiidae. In addition to morphological characters of the lineage, our molecular results support the separation of Borysthenidae as a valid, independent lineage deserving a family level within the Delphacoidea superfamily.

### Superfamily Delphacoidea Leach, 1815

Borysthenidae Emeljanov, 1989 **stat. rev.**

Borystheninae: Emeljanov, p. 94 [new subfamily].
Emeljanov, p. 106.
Holzinger et al., p. 124.
Liang, p. 809.
Ceotto and Bourgoin, p. 485.
Ceotto et al., p. 668.
Borysthenini: Luo et al., p. 15.
Borysthenini: Bucher et al., p. 11.

#### Diagnostic characters

Tegmen with clavus short, apex of clavus prolonged to tornus; tegmen without distinct pterostigma, pterostigmal area not thickened. Costal margin with veins of costal complex slightly shifted, with two types of sensory pits, larger and smaller with seta placed at anterior margin (CA); two types of secretory structures, with and without short seta of Pc+CP, separation of Pc+CP at connection with ScP(+RA1) absent; stem ScP+R, and terminal ScP(+RA1), with rounded secretory structures, arranged in rows; terminal ScP+(RA1) oblique, reaching the costal margin not fused with branching of Pc+CP of costal complex; tegmina with membrane distinctly widened, overlapping, covering the body.

Head with lateral ocellus and antennal fovea shifted to the front of the compound eye; compound eye vertical; subantennal expansion (subantennal carina: Emeljanov, 1989; subantennal process: Liang, 2005) on the gena, below the compound eye and antenna, above the level of frontoclypeal suture, provided with fine, elongate setae on outer margin and plaque sensillum. Metatibia with a row of small spiniform sensilla, apical row of teeth without diastema, the outermost not distinctly larger than preceding ones.

#### Content

Only genus *Borysthenes* Stål, 1866 (type species *Cixius dilectus* Walker, 1857), with species *Borysthenes acuminatus* Fennah, 1956; *Borysthenes certus* Muir, 1913; *Borysthenes cougus* Huang, 1995; *Borysthenes deflexus* Fennah, 1956; *Borysthenes dilectus* (Walker, 1857); *Borysthenes diversa* (Distant, 1906); *Borysthenes emarginatus* Fennah, 1956; *Borysthenes fascialatus* Muir, 1922; *Borysthenes fatalis* Emeljanov, 1989; *Borysthenes finitus* (Walker, 1857); *Borysthenes fusconotatus* (Melichar, 1914); *Borysthenes garambensis* Van Stalle, 1984; *Borysthenes hainanensis* Lyu & Webb, 2023; *Borysthenes incertus* Muir, 1913; *Borysthenes lacteus* Tsaur & Lee, 1987; *Borysthenes maculatus* (Matsumura, 1914); *Borysthenes magnus* Muir, 1913; *Borysthenes mambilensis* Van Stalle, 1984; *Borysthenes mlanjensis* Muir, 1923; *Borysthenes nambilensis* Van Stalle, 1984; *Borysthenes nicanor* Fennah, 1978; *Borysthenes ponomarenkoi* Emeljanov, 1989; *Borysthenes simulans Muir*, 1913; *Borysthenes strigipennis* Distant, 1911; *Borysthenes suknanicus* Distant, 1911).

### 3. Treatment and position of Achilixiidae Muir, 1923

The monophyletic family Achilixiidae Muir, 1923 is supported in our analysis, after examination of representatives of both known genera *Achilixius* Muir, 1923 and *Bebaiotes* Muir, 1924. The family was included in Achilidae as subfamilies Achilixiiinae Muir, 1923 and Bebaiotinae Emeljanov, 1991 (Emeljanov, 1991), but this statement was challenged by Liang (2001). Achilixiidae was treated as a separate family by Urban and Cryan (2007), Bartlett et al. (2018), and Viegas and Ale-Rocha (2024). The status and placement of Achilixiidae were recently discussed by Bucher et al. (2023), with *Bebaiotes* – the only analyzed genus – placed in the Achilidae-Derbidae clade. Our results, based on both subfamilies, firmly placed Achilixiidae outside of the Achilidae-Derbidae clade, and as a sister clade to the rest of the superfamily Fulgoroidea excluding Meenoploidea. However, we note that this placement may be subject to changes following more representative and comprehensive sampling in Achilidae and Derbidae.

### 4. Placement of the tribes Lyncidini and Amyclini in Dictyopharidae and the genus *Madagascaritia* in Fulgoridae

The tribe Lyncidini Schmidt, 1915 was originally placed in Fulgoridae in the subfamily Dictyopharinae (Schmidt, 1915). Muir (1923) transferred it to Dictyopharidae: Orgerinae. Then, Emeljanov (1969) placed it in Dictyopharidae, proposing the subfamily Lyncidinae with tribe Lyncidini (including also Capenini Emeljanov, 1969, Risiini Fennah, 1962 and Strongylodematini Fennah, 1962), and subsequently extracted Lyncidini and placed them in Fulgoridae: Lyncidinae: Lyncidini (Emeljanov, 1979), with only genus from Madagascar – *Lyncides* Stål, 1866. Our results support the decision of Emeljanov (1969) to place Lyncidini within Dictyopharidae.

Amyclini Metcalf, 1938 was first established as a subfamily in Fulgoridae (Metcalf, 1938), with two genera *Amycle* Stål, 1861 and *Scolopsella* Ball, 1905. Later, Metcalf (1947) proposed the tribe Amyclini Metcalf, 1938 and added the tribe Xosopharini Metcalf, 1947, which later was upgraded to a subfamily (Lallemand, 1959). Urban and Cryan (2009) found *Amycle* sister to the unplaced *Amerzanna*, which, together, were sister to Fulgoridae subfamilies Poiocerinae and Phenacinae. In contrast, our analyses strongly placed *R. brunnea* Van Duzee, 1929 inside Dictyopharidae, questioning the classification of the tribe Amyclini Metcalf, 1938 in Fulgoridae.

Additional evidence supporting the placements recovered by our analyses comes from the host plants. Species of the genus *Amycle* are associated with grasses (Order: Poales); grass feeding is more often associated with Dictyopharidae than with Fulgoridae, where it is rare (Emeljanov, 1979). *Rhabdocephala brunnea* Van Duzee, 1929 is associated with the grasses *Muhlenbergia porteri* in the family Poaceae, but was found also on Asteraceae *Baccharis sarothroides* (Wilson and Wheeler, 1992). Unfortunately, no host plant association is available for *Lyncides*.

The genus *Madagascaritia* Song et Liang, 2016 was initially classified within the tribe Aluntiini in the family Dictyopharidae (Song et al., 2016). The tribe and its genera are readily distinguishable based on their morphology, and the tribe itself constituted a discrete clade indicated by Song et al. (2016). To our knowledge, molecular data are only available for the genus *Madagascaritia*, which appears as the most basal offshoot within the Fulgoridae clade in our results. Therefore, we transfer this genus from the Dictyopharidae: Aluntiini to Fulgoridae, but leave the remaining Aluntiini clades in Dictyopharidae. The placement of other Aluntiini species and the tribe as a whole can only be accessed with a wider examination of this tribe and other fulgorids and dictyopharids. Taken together, these results indicate that the delineation between families Dictyopharidae and Fulgoridae and their inner classification is very challenging, which requires larger and broader sampling in the future.

### 5. Transfer of *‘Suva’ longipenna* Yang et Hu, 1985 from Meenoplidae to Derbidae

The species named *‘Suva’ longipenna* was described by Yang and Hu (1985: 24), based on the material from Tianmushan (Zhejiang, China). In the original description, the Authors stated it is very different in morphology and structure of male terminalia (Yang and Hu, 1985: fig. 3) from other species of the genus *Suva* Kirkaldy, 1906. In the same paper, another species ascribed to the genus *Suva* was proposed as *‘Suva’ flavimaculata* Yang et Hu, 1985 (Yang and Hu, 1985: 25; fig. 4). Bourgoin (1997) excluded both species from Meenoplidae and transferred them to Derbidae. Wang et al. (2023) found *‘Suva’ longipenna* clustered with *Diostrombus politus* Uhler, 1896 (Derbidae: Otiocerinae: Zoraidini: Lyddina), which corroborates the placement of this species in Derbidae. Nevertheless, as of July 2024, both names appear as valid in Meenoplidae in the FLOW database (Bourgoin, 2024). Here we propose to place *‘Suva’ longipenna* Yang et Hu, 1985 as species in Derbidae: Otiocerinae, and *‘Suva’ flavimaculata* Yang et Hu, 1985 as *incertae sedis* species in Derbidae.

### 6. Nomenclatural note on GenBank data – *‘Zanna’ intricata* Walker, 1858

The GenBank database lists the genome assembly ASM1001600v3 (GCA_010016005.3) as representing *Zanna intricata* Walker, 1858 (Fulgoridae, Zanninae), which is a species known from a few localities on the southeastern coast of the African continent. However, this classification is questionable given the clear phylogenetic placement of this species within the genus *Pyrops* Spinola, 1839. This led us to conclude that the specimen was misidentified and most probably represents *Pyrops intricatus* (Walker, 1857), known from Indonesia (Borneo, Kalimantan) and Malaysia (Sabah, Sarawak). *Pyrops intricatus* (Walker, 1857) was placed in Fulgoridae *incertae sedis* in the FLOW database (Bourgoin, 2024), but proposed as representing the *pyrorhynchus* group of the genus *Pyrops* by Constant (2015).

## Discussion

### Phylogenomic datasets yielded the most robust phylogenetic hypothesis of planthoppers so far

The inclusion of information from 1164 nuclear markers inferred trees with strong supports compared to the ones based on only mitogenomes, few markers, or morphology, providing many new insights into the relationship among worldwide planthopper species (Figs. 1 & S4). Our best trees not only confirmed well-accepted relationships but also resolved controversies and proposed strongly supported, novel placements for many clades, some of which were recovered for the first time. Taxonomic decisions based on these data include the establishment of a new family Borysthenidae **stat. rev.** within Delphacoidea, redefinition of the family Cixiidae, establishment of a new superfamily Meenoploidea **superfam. nov.** with redefined Kinnaridae **stat. rev.** and Meenoplidae **stat. rev**., and confirmation of the monophyletic family Achilixiidae outside the Achilidae-Derbidae clade. Furthermore, by increasing our sampling with additional mitogenomes from GenBank, we show detailed relationships among tribes and subfamilies within each family (Fig. 1). We discuss the findings within each family in detail in the following sections.

### The superfamily Delphacoidea and the content of Cixiidae

The sister relationship between Delphacidae and Cixiidae has been recovered by all published studies based on either a single or multiple genetic markers (Bucher et al., 2023; Song and Liang, 2013; Urban and Cryan, 2007; Wang et al., 2023; Yeh et al., 2005). Within Cixiidae, the placement of *Borysthenes* outside the sister clade of Delphacidae and Cixiinae was not recovered by previous phylogenetic hypotheses of cixiids based on few markers (Bucher et al., 2023; Ceotto et al., 2008). Wang et al. (2023) recovered a similar paraphyly of Cixiidae in one analysis but not in the other based on complete mitogenomes, which probably resulted from poor sampling (only five cixiids were included). The molecular placement of this clade and distinct morphology from other cixiids justify the proposal of an independent family Borysthenidae **stat. rev.** within the Delphacoidea superfamily (see above “Taxonomic decisions” section). On the other hand, the internal classification of Cixiidae, including the concept and content of suprageneric taxa, remains fragile due to the lack of molecular data in taxa showing ancestral or peculiar morphological features (e.g., Mnemosynini, Brixiini, Brixidiini, Andini) or with complex internal taxonomic content (e.g., Cixiini and related tribes). Future sampling in these taxa will be crucial to fully understand the relationships among cixiids.

The relationships among delphacids are generally well-understood. Studies based on few markers recovered the same relationships among Delphacidae subfamilies (Asiracinae, Vizcayinae, Kelisiinae, Stenocraninae, and Delphacinae) as we did here (Huang et al., 2020; Urban et al., 2010; Wang et al., 2023). Bucher et al. (2023) separated Asiracinae from the monophyly of the remaining subfamilies based on six genes, but the support was weak. Wider geographical sampling for Asiracinae tribes, especially the New World species in Ugyopini, may change the understanding of phylogeny within this clade. Likewise, additional sampling from geographically underrepresented areas (e.g., Subsaharean Africa and southern South America), habitats, and host plants may change the topology of the species-rich and agriculturally significant subfamily Delphacinae, which we reconstructed as comprising three well-separated tribes.

### The status of Kinnaridae and Meenoplidae

The placement of Kinnaridae and Meenoplidae as a monophyletic clade that branches off early in Fulgoroidea has not been much doubted since the same topology was repeatedly recovered by previously published trees based on few samples and genes (Bucher et al., 2023; Song and Liang, 2013; Urban and Cryan, 2007; Wang et al., 2023). Within each family, the monophyly was recovered consistently for Meenoplidae, but not for Kinnaridae. Bucher et al. (2023) found the paraphyly of Kinnaridae by placing *Kinara ochracea* (subfamily Kinnarinae) as sister to the monophyletic clade formed by the rest of kinnarids – all from the subfamily Prosotropinae – and meenoplids. This topology is similar to ours when we included the unrecognized kinnarid from Wang et al. (2023), indicating that this species likely represents the genus *Kinnara*. These results support the proposal of the new superfamily Meenoploidea **superfam. nov.** and the reorganization of families Kinnaridae and Meenoplidae (see “Taxonomic decisions” section). In the future, more sampling in the subfamilies and tribes is needed to understand the morphological diversity and relationships among Kinnaridae and Meenoplidae lineages.

### Achilixiidae as a monophyletic family separated from Achilidae and Derbidae

The placement of the family Achilixiidae has been controversial for decades. Published phylogenetic hypotheses consistently placed achilixiids close to the families Achilidae or Derbidae, which were often recovered paraphyletic, with weak support and little agreement among hypotheses (Bucher et al., 2023; Song and Liang, 2013; Urban and Cryan, 2007). Our analyses clearly indicate that Achilixiidae is a distinct and well-established monophyletic family outside of Achilidae and Derbidae, a classification that had previously been questioned based on inconclusive phylogenetic analyses (Bartlett et al., 2018; Brysz and Szwedo, 2019; Emeljanov, 1991; Viegas, 2023).

The sister relationship between Achilidae and Derbidae and their earlier branch-off in Fulgoroidea than Fulgoridae and Dictyopharidae was previously recovered by Bucher et al. (2023) based on six genes and a larger sample size. Skinner et al. (2020) found a similar topology based on transcriptomes, although they failed to recover the sister relationship between Achilidae and Derbidae, likely due to a limited sampling. On the other hand, the trees inferred from mitogenomes in our analyses and by Wang et al. (2023) tended to recover the Fulgoridae-Dictyopharidae group as an earlier branch-off than Achilidae and Derbidae, with weak support for this alternative topology.

### Gondwanaland as the potential cradle of fulgorids and dictyopharids

The sister relationship we found between Fulgoridae and Dictyopharidae has been repeatedly inferred from molecular data (Bucher et al., 2023; Song and Liang, 2013; Urban and Cryan, 2007; Wang et al., 2023). However, there has been ongoing debate about the classification of clades within these two families, with many groups repeatedly transferred between them (Song et al., 2018, 2016). Our phylogenies suggested the misclassification of several species, including *Lyncides* sp., *Madagascaritia* sp., and *R. brunnea,* all placed inside the family to which they are not currently classified. A relatively well-known case comes from the genus *Zanna*. Urban and Cryan (2009) argued that *Zanna* belongs to Dictyopharidae based on the phylogeny inferred from five genes. On the contrary, we firmly placed *Z. robusticephalica* within Fulgoridae, in agreement with recent phylogenies based on a larger sample size or mitogenomes (Bucher et al., 2023; Wang et al., 2023). These misclassifications suggested that the taxonomy within Fulgoridae and Dictyopharidae needs to be revised.

The placement of Madagascar species *Lyncides* sp. and *Madagascaritia* sp. as the most anciently diversifying lineages in their respective families suggests a Gondwana origin of Fulgoridae and Dictyopharidae. An estimated root age of ∼140 Ma for these two families is consistent with the timing of the break-up of Gondwana, indicating that they diversified contemporaneously as the Gondwanaland fractured (Matthews et al., 2016). However, further characterization of the diversification of fulgorids and dictyopharids requires additional sampling. Urban and Cryan (2009) found a monophyletic clade comprising all sampled New World fulgorids and suggested two alternative biogeographic hypotheses for the diversification of New World and Old World lineages based on their relative placements on the tree. Unfortunately, our analysis did not include any fulgorids from the New World. In Dictyopharidae, our results showed mixed relationships among New World and Old World lineages with no clear separation between them. Future samplings from under-represented groups and regions (e.g., Africa, Madagascar) are needed to fully understand the origin and diversification of these two families.

### Tropiduchidae

The monophyly of Tropiduchidae and its sister relationship to the remaining families of Fulgoroidea has been repeatedly recovered by previous studies based on broad sampling (11 species; Bucher et al., 2023), complete mitogenomes (three species; Wang et al., 2023), and transcriptomics (one species; Skinner et al., 2020). However, the sampling remains scarce within Tropiduchidae. We only included less than one-third of established tribes (five out of ∼25) in Tropiduchidae, missing some of those with distinct morphology and ecological adaptations (Fennah, 1982; Gnezdilov et al., 2016; Gnezdilov and Bourgoin, 2015; Stroiński et al., 2022; Wang et al., 2014). Their future characterization of species from these missing tribes may challenge the monophyly of this family (Wang et al., 2017).

### Tettigometridae, Caliscelidae, Lophopidae, and Eurybrachidae

There has been ongoing debate about whether the family Tettigometridae is an ancient planthopper lineage with primitive morphological traits and lifestyle or a young family (Bourgoin et al., 1997). Most published trees placed Tettigometridae as a young family branching off late in Fulgoroidea (Bourgoin et al., 1997; Song and Liang, 2013; Urban and Cryan, 2007). In contrast, Bucher et al. (2023), based on six markers and a broad sampling, placed Tettigometridae as a clade that branched off early in Fulgoroidea. Our analyses strongly support the former conclusion – a late branch-off of Tettigometridae in Fulgoroidea. We suspect that Bucher et al.’s unusual placement of Tettigometridae might have resulted from long-branch attraction due to short alignments and a broad sampling. In addition, we recovered the sister relationship between Tettigometridae and Caliscelidae, previously reported by Skinner et al. (2020) and Song and Liang (2013).

The sister relationship of the Lophopidae and Eurybrachidae that we found was recovered repeatedly by previous studies (Bucher et al., 2023; Urban and Cryan, 2009; Wang et al., 2023). Additionally, we recovered them as a sister clade to the Tettigometridae-Caliscelidae clade. Wang et al. (2023) found a similar topology of Lophopidae, Eurybrachidae, and Caliscelidae based on complete mitogenomes but did not include any tettigometrids in their phylogeny. Skinner et al. (2020) recovered a different topology where the Tettigometridae-Caliscelidae clade branched off earlier than Eurybrachidae, despite that only one or two species of each family were sampled and no lophopids were included. Bucher et al. (2023) recovered a similar four-family monophyletic group but placed Acanaloniidae, instead of Tettigometridae, sister to Calisceliidae.

### Broader sampling and genomic investigation are needed to resolve the phylogeny of the remaining families

The relationships among relatively young families Ricaniidae, Issidae, Nogodinidae, Acanaloniidae, and Flatidae were weakly supported and were represented by short internal branches on our phylogeny (Figs. 1 & S4), indicating a history of rapid diversification within a relatively short timeframe. Our time-calibrated phylogeny suggested that these families appeared in a period of 20 million years in the Cretaceous, from ∼135 Ma when their common ancestral lineage started diversifying to ∼115 Ma when all families were present. This period saw the flourishing of flowering plants on earth, which likely drove the rapid diversification and species turnover among insects (Peris and Condamine, 2024).

Such rapid radiations make phylogenetic inference challenging and susceptible to random errors (Simion et al., 2020). These effects were clearly shown by early phylogenetic efforts based on few molecular markers when little agreement on the topology was found (Bucher et al., 2023; Song and Liang, 2013; Urban and Cryan, 2007; Wang et al., 2023).

Based on much greater amounts of data, our analyses provided better-supported topologies compared to the published studies. Nogodinidae is known for its paraphyly, with historically repeated species transfer between Nogodinidae, Ricaniidae, and Issidae (Song and Liang, 2013; Urban and Cryan, 2007; Wang et al., 2016). In agreement with these studies, the nogodinids we characterized grouped into at least three distinct clades (with low branch support preventing firm conclusions about their count), recovered as sisters to Ricaniidae, Issidae, and the Acanaloniidae-Flatidae clades. These results further question the family status of Nogodinidae and call for an urgent revision of this group using integrative molecular and morphological methods based on broad sampling that includes the type genus *Nogodina*, morphologically distinctive and geographically isolated groups (e.g., Australian lineages), and good representations of related families.

We confirmed the monophyly of other families in this clade: Ricaniidae, Issidae, Acanaloniidae, and Flatidae. Additionally, we placed Acanaloniidae as sister to Flatidae, in agreement with Skinner et al. (2020) based on transcriptomics. Within Issidae, our analyses agreed with previously inferred phylogenies, recovering the monophyly of the subfamilies Hysteropterinae and Thioniinae, the paraphyly of the subfamily Hemisphaeriinae, and the reciprocal monophyly of tribes Hemisphaeriini and Parahiraciini within Hemisphaeriinae (Bucher et al., 2023; Gnezdilov et al., 2022).

### New fossils and genomic data yielded the oldest estimate of the origin of Fulgoromorpha

Our analyses, based on genomic data for a taxonomically comprehensive planthopper collection and 30 carefully selected fossils, pushed back the origin of Fulgoromorpha relative to previous studies. We estimated an age of ∼263 Ma for the crown Fulgoromorpha with our age constraints and ∼332 Ma with alternative constraints. Johnson et al. (2018) placed the origin of Fulgoromorpha at the upper Triassic of ∼206 Ma (95% CI: 178 - 241 Ma) based on transcriptomics, but only three planthopper fossils and 13 planthopper species from nine families were included. Wang et al. (2023) performed simple analyses with two fossils on a mitogenome-based phylogeny, estimating a Lower Jurassic origin of ∼193 Ma for Fulgoromorpha. In the same year, Bucher et al. (2023) used ten fossils on the tree of 531 species inferred by six genes, placing the origin of extant Fulgoromorpha at ∼238 Ma with a wide 95% CI (199.6–352.9 Ma). We notice the gap between the age of the oldest fossil (∼125 Ma) representing the crown group and our estimates. Schachat et al. (2022), by comprehensively investigating and comparing fossil records between insect groups, argued how such gaps might represent analytical artifacts resulting from taxon sampling, fossil choices, and early history of species diversification. On the other hand, the current fossil record is far from complete. The fossilization processes can be biased, depending on the conditions in past environments, the size of the insects, their paleoecology and palaeobehavior, and numerous other taphonomic reasons (Karr and Clapham, 2015; Martínez-Delclòs et al., 2004). The discovery of older fossils might push the age of certain taxa deeper in evolutionary history. The newly proposed placement of fossils in the phylogeny can also significantly change our interpretation of species history. Bourgoin and Szwedo (2022, 2023) proposed that the oldest probable stem-group hemipteran *Aviorrhyncha magnifica*, dated back to Moscovian, Carboniferous (∼310 Ma), should be assigned to Fulgoromorpha *incertae sedis*. The dynamics in fossil research and the usage of different models and methods are likely to change our understanding of the history of Fulgoromorpha in the future.

## Conclusion

Our study highlights that nuclear genome-level data offers superior phylogenetic resolution, indicating that the combination of genomic and morphological approaches is essential for the robust placement of newly described clades and the revisions of currently proposed relationships among known clades. Within planthoppers, however, the sequencing speed is far behind the pace of species discovery and sampling, with many new species described but little molecular data, especially genome-level data, generated. With the rapidly decreasing sequencing costs, we hope that the sequence alignments and bioinformatic pipeline presented here will encourage and facilitate future studies to use genomic data and phylogenomic tools in the systematic research on planthoppers.

At the same time, the power and robustness of phylogenomic analyses are often limited by insufficient sampling. The inclusion of 285 planthopper species here – the second largest sampling after 531 species in Bucher et al. (2023) – is still relatively poor, given a total of >14000 described species. The scarcity of molecular data from morphologically and ecologically diverse families, such as Tropiduchidae, Nogodinidae, and Flatidae, hinders conclusions about their family status and the relationships among subfamilies or tribes within. The lack of sampling from geographically remote and under-represented regions (e.g., Sub-Saharan Africa, South America, Australia) and poorly characterized niches (e.g., tree canopies, underground environments, caves) leaves a great diversity unexplored and a great number of newly described taxa unplaced on the phylogeny. A great example comes from the two South African families Hypochthonellidae and Gengidae, known only from a few collected specimens of a few described species. With little morphological description and no published molecular data, their placement on planthopper phylogeny is still a mystery. In contrast, as an example from a different hemipteran clade, the molecular characterization of an inconspicuous small cicada, *Derotettix mendosensis*, captured in the degraded salt-plain habitats in arid regions of central Argentina, has led to the description of a new subfamily that is sister to all other living cicadas, bringing about new hypotheses of Cicadidae origins (Simon et al., 2019). Future sampling from these understudied taxa, regions, and habitats, combined with genomic and morphological tools, is likely to challenge the current topology of planthopper phylogeny and change our view of evolutionary history and taxonomic assignment of planthoppers.

Additionally, we view the accurate reconstruction of species relationships as not only the goal of phylogenomic research but also a way to provide the essential framework for the understanding of biotic interactions and symbioses between planthoppers and other organisms. Given their roles as carriers of plant pathogens, as pests of agricultural plants, as food resources of other animals, and as hosts of diverse microbial symbionts, the phylogeny will allow future studies to trace the co-evolutionary history between planthoppers and other organisms, providing insights into pathogen transmission, pest control, and genomic and functional evolution of symbionts.

## Data availability

The gene alignments and output of all phylogenetic analyses, including raw tree files from IQ-TREE2 and the log files from BEAST runs, will be available on Dryad (DOI: 10.5061/dryad.3ffbg79sc).

## Acknowledgments

We thank Chris Dietrich, Charles Bartlett, and Brian Fisher for sharing specimens from their collections. We thank Matthew P. Greenwood for introducing the ortholog assembly tool HybPiper to us. We gratefully acknowledge Polish high-performance computing infrastructure PLGrid (HPC Centers: WCSS) for providing computer facilities and support within computational grant no. PLG/2023/016612. This project was also supported by the Polish National Science Centre grants 2017/26/D/NZ8/00799 (to A.M.) and 2018/30/E/NZ8/00880 (to P.Ł.) as well as the Polish National Agency for Academic Exchange grant PPN/PPO/2018/1/00015 (to P.Ł.), and Jagiellonian University Visibility and Mobility Module grant WSPR.WSDNSP.2.5.2022.4 (to J.D.).

## Author contributions

**Junchen Deng**: Writing – Original Draft (lead); Conceptualization (equal); Data Curation (lead); Formal Analysis (lead); Visualization (lead); Writing – Review & Editing (equal); Funding Acquisition (equal)

**Adam Stroiński**: Writing – Original Draft (equal); Conceptualization (equal); Resources (lead); Writing – Review & Editing (equal)

**Jacek Szwedo**: Writing – Original Draft (equal); Conceptualization (equal); Formal Analysis (supporting); Resources (lead); Writing – Review & Editing (equal)

**Hamid Reza Ghanavi**: Formal Analysis (supporting); Writing – Review & Editing (equal)

**Etka Yapar**: Formal Analysis (supporting); Writing – Review & Editing (equal)

**Diego Castillo Franco**: Data Curation (supporting); Writing – Review & Editing (equal)

**Monika Prus-Frankowska**: Resources (equal); Writing – Review & Editing (equal)

**Anna Michalik**: Resources (equal); Writing – Review & Editing (equal); Funding Acquisition (equal)

**NiklasWahlberg**: Formal Analysis (supporting); Writing – Review & Editing (equal)

**Piotr Łukasik**: Writing – Original Draft (supporting); Conceptualization (equal); Writing – Review & Editing (equal); Funding Acquisition (lead); Supervision (lead)

## Supporting information legends

**Figure. S1** Heat maps generated by the tests for among lineage compositional heterogeneity.

**Figure S2.** The best Maximum-Likelihood tree based on the concatenated nucleotide dataset of 13 mitochondrial protein-coding genes (PCGs) of 149 planthopper species newly sequenced in this study.

**Figure S3.** The best Maximum-Likelihood tree based on the concatenated nucleotide dataset of 13 mitochondrial PCGs of 282 planthopper species.

**Figure S4.** The best Maximum-Likelihood tree based on the concatenated nucleotide dataset of 1164 nuclear markers of 149 planthopper species newly sequenced in this study.

**Figure S5.** The best Maximum-Likelihood tree based on the concatenated nucleotide dataset of 1164 nuclear markers and 13 mitochondrial PCGs of 246 planthopper species. Forty species from Wang et al. (2023) are excluded from this phylogeny.

**Figure S6.** The species tree of 149 species inferred from 1164 nuclear gene trees by ASTRAL.

**Figure S7.** The dated phylogeny shown in Figure 2 with the mean ages of each node shown.

**Figure S8.** A similar dated phylogeny to Figure S7 under the alternative set of age constraints based on the published dated tree from Johnson et al. (2018).

**Table S1**. Species list

**Table S2**. List of fossils and age constraints

**Table S3**. The number of assembled nuclear markers for each sample (1164 markers in total)

**Table S4**. The credits to photos taken from iNaturalist.org that were used for the illustration purpose

## References

Anisimova, M., Gascuel, O., 2006. Approximate Likelihood-Ratio Test for Branches: A Fast, Accurate, and Powerful Alternative. Syst. Biol. 55, 539–552. 10.1080/10635150600755453

Anisimova, M., Gil, M., Dufayard, J.-F.O., Dessimoz, C., Gascuel, O., 2011. Survey of Branch Support Methods Demonstrates Accuracy, Power, and Robustness of Fast Likelihood-based Approximation Schemes. Syst. Biol. 60, 685–699. 10.1093/sysbio/syr041

Bartlett, C.R., Deitz, L.L., Dmitriev, D.A., Sanborn, A.F., Soulier-Perkins, A., Wallace, M.S., 2018. The Diversity of the True Hoppers (Hemiptera: Auchenorrhyncha), in: Insect Biodiversity: Science and Society, II. Wiley-Blackwell, Oxford, Oxford, pp. 501–590. 10.1002/9781118945582.ch19

Bennett, G.M., Moran, N.A., 2013. Small, Smaller, Smallest: The Origins and Evolution of Ancient Dual Symbioses in a Phloem-Feeding Insect. Genome Biol. Evol. 5, 1675–1688. 10.1093/gbe/evt118

Bouckaert, R., Vaughan, T.G., 2019. BEAST 2.5: An advanced software platform for Bayesian evolutionary analysis. PLoS Comput. Biol. 15, e1006650. 10.1371/journal.pcbi.1006650

Bourgoin, T., 2024. FLOW (Fulgoromorpha Lists on The Web): a world knowledge base dedicated to Fulgoromorpha. Version 8, updated 2024-01-04 [WWW Document]. URL https://www.hemiptera-databases.org/flow/

Bourgoin, T., 1997. The Meenoplidae (Hemiptera, Fulgoromorpha) of New Caledonia, with a revision of the genus Eponisia Matsumura, 1914, and new morphological data on forewing venation and wax plate areas. Memoires Mus. Natl. D’Histoire Nat. 171, 197–249.

Bourgoin, T., 1993. Female Genitalia in Hemiptera Fulgoromorpha, Morphological and Phylogenetic Data. Ann. Société Entomol. Fr. NS 29, 225–244. 10.1080/21686351.1993.12277686

Bourgoin, T., Gjonov, I., Lapeva-Gjonova, A., Roger, S., Constant, J., Kunz, G., Wilson, M.R., 2023. When Cockroaches Replace Ants in Trophobiosis: A New Major Life-Trait Pattern of Hemiptera Planthoppers Behaviour Disclosed When Synthesizing Photographic Data. Diversity 15, 356. 10.3390/d15030356

Bourgoin, T., Steffen-Campbell, J.D., Campbell, B.C., 1997. Molecular Phylogeny of Fulgoromorpha (Insecta, Hemiptera, Archaeorrhyncha). The Enigmatic Tettigometridae: Evolutionary Affiliations and Historical Biogeography. Cladistics 13, 207–224. 10.1111/j.1096-0031.1997.tb00316.x

Bourgoin, T., Szwedo, J., 2023. Toward a new classification of planthoppers Hemiptera Fulgoromorpha: 2. Higher taxa, their names and their composition. Zootaxa 5297, 562–568. 10.11646/zootaxa.5297.4.5

Bourgoin, T., Szwedo, J., 2022. Toward a New Classification of Planthoppers Hemiptera Fulgoromorpha: 1. What Do Fulgoridiidae Really Cover? Ann. Zool. 72, 951–962. 10.3161/00034541ANZ2022.72.4.011

Bradley, R.K., Roberts, A., Smoot, M., Juvekar, S., Do, J., Dewey, C., Holmes, I., Pachter, L., 2009. Fast Statistical Alignment. PLoS Comput. Biol. 5, e1000392. 10.1371/journal.pcbi.1000392

Brysz, A.M., Szwedo, J., 2019. Jeweled Achilidae – a new look at their systematics and relation to other Fulgoroidea (Hemiptera). Monogr. Up. Silesian Mus. 10, 93–130. 10.5281/zenodo.3600279

Bucher, M., Condamine, F.L., Luo, Y., Wang, M., Bourgoin, T., 2023. Phylogeny and diversification of planthoppers (Hemiptera Fulgoromorpha) based on a comprehensive molecular dataset and large taxon sampling. Mol. Phylogenet. Evol. 107862. 10.1016/j.ympev.2023.107862

Bucher, M., Gignoux, G., Szwedo, J., Bourgoin, T., 2024. Time-traveling through fossil planthopper tegmina in the Paleozoic and Mesozoic eras (Insecta: Hemiptera: Fulgoromorpha). Palaeoentomology 7, 1–67. 10.11646/palaeoentomology.7.1.1

Buchfink, B., Reuter, K., Drost, H.-G., 2021. Sensitive protein alignments at tree-of-life scale using DIAMOND. Nat. Methods 18, 366–368. 10.1038/s41592-021-01101-x

Capella-Gutierrez, S., Silla-Martinez, J.M., Gabaldon, T., 2009. trimAl: a tool for automated alignment trimming in large-scale phylogenetic analyses. Bioinformatics 25, 1972–1973. 10.1093/bioinformatics/btp348

Ceotto, P., Bourgoin, T., 2008. Insights into the phylogenetic relationships within Cixiidae (Hemiptera: Fulgoromorpha): cladistic analysis of a morphological dataset. Syst. Entomol. 33, 484–500. 10.1111/j.1365-3113.2008.00426.x

Ceotto, P., Kergoat, G.J., Rasplus, J.-Y., Bourgoin, T., 2008. Molecular phylogenetics of cixiid planthoppers (Hemiptera: Fulgoromorpha): New insights from combined analyses of mitochondrial and nuclear genes. Mol. Phylogenet. Evol. 48, 667–678. 10.1016/j.ympev.2008.04.026

Chan, P.P., Lin, B.Y., Mak, A.J., Lowe, T.M., 2021. tRNAscan-SE 2.0: improved detection and functional classification of transfer RNA genes 49, 9077–9096. 10.1093/nar/gkab688

Chernomor, O., von Haeseler, A., Minh, B.Q., 2016. Terrace Aware Data Structure for Phylogenomic Inference from Supermatrices. Syst. Biol. 65, 997–1008. 10.1093/sysbio/syw037

Constant, J., 2015. Review of the effusus group of the lanternfly genus Pyrops Spinola, 1839, with one new species and notes on trophobiosis (Hemiptera: Fulgoromorpha: Fulgoridae). Eur. J. Taxon. 128, 1–23. 10.5852/ejt.2015.128

Davranoglou, L.-R., Hartung, V., 2024. Moss bugs shed light on the evolution of complex bioacoustic systems. PLOS ONE 19, e0298174. 10.1371/journal.pone.0298174

Dejean, A., Bourgoin, T., Orivel, J., 2000. Ant Defense of *Euphyonarthex phyllostoma* (Homoptera: Tettigometridae) during Trophobiotic Associations. Biotropica 32, 112–119.

Deng, J., Bennett, G.M., Franco, D.C., Prus-Frankowska, M., Stroiński, A., Michalik, A., Łukasik, P., 2023. Genome Comparison Reveals Inversions and Alternative Evolutionary History of Nutritional Endosymbionts in Planthoppers (Hemiptera: Fulgoromorpha). Genome Biol. Evol. 15, evad120. 10.1093/gbe/evad120

Dietrich, C.H., 2009. Chapter 15 - Auchenorrhyncha: (Cicadas, Spittlebugs, Leafhoppers, Treehoppers, and Planthoppers), in: Resh, V.H., Cardé, R.T. (Eds.), Encyclopedia of Insects (Second Edition). Academic Press, San Diego, pp. 56–64. 10.1016/B978-0-12-374144-8.00015-1

Douglas, J., Zhang, R., Bouckaert, R., 2021. Adaptive dating and fast proposals: Revisiting the phylogenetic relaxed clock model. PLoS Comput. Biol. 17, e1008322. 10.1371/journal.pcbi.1008322

Eddy, S.R., 2011. Accelerated Profile HMM Searches. PLoS Comput. Biol. 7, e1002195. 10.1371/journal.pcbi.1002195

Emeljanov, A.F., 2002. Contribution to classification and phylogeny of the family Cixiidae (Hemiptera, Fulgoromorpha). Denisia 4, 103–112.

Emeljanov, A.F., 1991. K voprosu ob obeme i podrazdeleniyakh sem. Achlidae (Homoptera, Cicadina). Entomol. Obozr. 70, 373–393.

Emeljanov, A.F., 1990. An attempt of construction of the phylogenetic tree of the planthoppers (Homoptera, Cicadina). Ėntomologicheskoe Obozr. 69, 353–356.

Emeljanov, A.F., 1989. To the problem of division of the family Cixiidae (Homoptera, Cicadina). Entomol. Obozr. 68, 93–106 [In Russian].

Emeljanov, A.F., 1979. The problem of family distinction between the Fulgoridae and the Dictyopharidae (Homoptera, Auchenorrhyncha). Tr. Zool. Instituta 82, 3–22 [In Russian].

Emeljanov, A.F., 1969. Reclassification of Palearctic planthoppers of the subfamily Orgeriinae (Homoptera, Dictyopharidae). Entomol. Obozr. 48, 189–198 [In Russian].

Fennah, R.G., 1982. A tribal classification of the Tropiduchidae (Homoptera: Fulgoroidea), with the description of a new species on tea in Malaysia. Bull. Entomol. Res. 72, 631–643.

Ghanavi, H.R., Twort, V., Hartman, T.J., Zahiri, R., Wahlberg, N., 2022. The (non) accuracy of mitochondrial genomes for family-level phylogenetics in Erebidae (Lepidoptera). Zool. Scr. 51, 695–707. 10.1111/zsc.12559

Gnezdilov, V.M., Bartlett, C.R., Bourgoin, T., 2016. A new tribe of Tropiduchidae (Hemiptera, Fulgoroidea) with revision of the genus Buca Walker and description of asymmetric hind leg spinulation. Fla. Entomol. 99, 406–416.

Gnezdilov, V.M., Bourgoin, T., 2015. New genera and new species of the tribe Elicini (Hemiptera: Fulgoroidea: Tropiduchidae) with key to Tropiduchid genera known from Madagascar. Ann. Zool. 65, 599–618.

Gnezdilov, V.M., Konstantinov, F.V., Namyatova, A.A., 2022. From modern to classic: Classification of the planthopper family Issidae (Hemiptera, Auchenorrhyncha, Fulgoroidea) derived from a total-evidence phylogeny. Syst. Entomol. 47, 551–568. 10.1111/syen.12546

Heong, K.L., Cheng, J., Escalada, M.M. (Eds.), 2015. Rice Planthoppers: Ecology, Management, Socio Economics and Policy, 1st ed. 2015. ed. Springer Netherlands : Imprint: Springer, Dordrecht. 10.1007/978-94-017-9535-7

Hoang, D.T., Chernomor, O., 2017. UFBoot2: Improving the Ultrafast Bootstrap Approximation. Syst. Biol. 35, 518–522. 10.1093/molbev/msx281

Holzinger, W.E., Emeljanov, A.F., Kammerlander, I., 2002. The family Cixiidae Spinola 1839 (Hemiptera: Fulgoromorpha)—a review. Denisia 4, 113–138.

Huang, Y.-X., Ren, F.-J., Bartlett, C.R., Wei, Y.-S., Qin, D.-Z., 2020. Contribution to the mitogenome diversity in Delphacinae: Phylogenetic and ecological implications. Genomics 112, 1363–1370. 10.1016/j.ygeno.2019.08.005

Jermiin, L.S., Ho, S.Y.W., Ababneh, F., Robinson, J., Larkum, A.W.D., 2004. The Biasing Effect of Compositional Heterogeneity on Phylogenetic Estimates May be Underestimated. Syst. Biol. 53, 638–643. 10.1080/10635150490468648

Johnson, K.P., Dietrich, C.H., Friedrich, F., Beutel, R.G., Wipfler, B., Peters, R.S., Allen, J.M., Petersen, M., Donath, A., Walden, K.K.O., Kozlov, A.M., Podsiadlowski, L., Mayer, C., Meusemann, K., Vasilikopoulos, A., Waterhouse, R.M., Cameron, S.L., Weirauch, C., Swanson, D.R., Percy, D.M., Hardy, N.B., Terry, I., Liu, S., Zhou, X., Misof, B., Robertson, H.M., Yoshizawa, K., 2018. Phylogenomics and the evolution of hemipteroid insects. Proc. Natl. Acad. Sci. 115, 12775– 12780. 10.1073/pnas.1815820115

Johnson, M.G., Gardner, E.M., Liu, Y., Medina, R., Goffinet, B., Shaw, A.J., Zerega, N.J.C., Wickett, N.J., 2016. HybPiper: Extracting coding sequence and introns for phylogenetics from high-throughput sequencing reads using target enrichment. Appl. Plant Sci. 4, 1600016. 10.3732/apps.1600016

Kalyaanamoorthy, S., Minh, B.Q., Wong, T.K.F., von Haeseler, A., Jermiin, L.S., 2017. ModelFinder: fast model selection for accurate phylogenetic estimates. Nat. Methods 14, 587–589. 10.1038/nmeth.4285

Karr, J.A., Clapham, M.E., 2015. Taphonomic biases in the insect fossil record: shifts in articulation over geologic time. Paleobiology 41, 16–32. 10.1017/pab.2014.3

Katoh, K., Standley, D.M., 2013. MAFFT Multiple Sequence Alignment Software Version 7: Improvements in Performance and Usability. Mol. Biol. Evol. 30, 772–780. 10.1093/molbev/mst010

Lallemand, V., 1959. Revision des especes africaines de la famille Fulgoridae (super-famille Fulgoroides-sous-ordre des Homopteres). Publicaçoes Cult. Cia. Diam. Angola Diamang Lisb. 41, 37–123.

Lehouck, V.S., Bonte, D.B., Dekoninck, W., Maelfait, J.-P.E., 2004. Trophobiotic relationships between ants (Hymenoptera: Formicidae) and Tettigometridae (Hemiptera: Fulgoromorpha) in the grey dunes of Belgium. Eur. J. Entomol. 101, 547–553. 10.14411/eje.2004.078

Li, D., Luo, R., Liu, C.-M., Leung, C.-M., Ting, H.-F., Sadakane, K., Yamashita, H., Lam, T.-W., 2016. MEGAHIT v1.0: A fast and scalable metagenome assembler driven by advanced methodologies and community practices. Methods 102, 3–11. 10.1016/j.ymeth.2016.02.020

Liang, A.-P., 2005. A new structure on the subantennal process of *Borysthenes* species (Hemiptera: Fulgoromorpha: Cixiidae: Borystheninae). Proc. Biol. Soc. Wash. 118, 809–814. 10.2988/0006-324X(2005)118[809:ANSOTS]2.0.CO;2

Liang, A.-P., 2001. Morphology of antennal sensilla in Achilixius sandakanensis Muir (Hemiptera: Fulgoromorpha: Achilixiidae) with comments on the phylogenetic position of the Achilixiidae. Raffles Bull. Zool. 49, 221–226.

Liang, A.-P., Yang, J.-T., 2001. The external male genitalia of Hemiptera (Homoptera — Heteroptera). Orient. Insects 35, 66. 10.1080/00305316.2001.10417287

Łukasik, P., Chong, R.A., Nazario, K., Matsuura, Y., Bublitz, D.A.C., Campbell, M.A., Meyer, M.C., Van Leuven, J.T., Pessacq, P., Veloso, C., Simon, C., McCutcheon, J.P., 2019. One Hundred Mitochondrial Genomes of Cicadas. J. Hered. 110, 247–256. 10.1093/jhered/esy068

Luo, Y., Bourgoin, T., Szwedo, J., Feng, J.-N., 2021. Acrotiarini trib. nov., in the Cixiidae (Insecta, Hemiptera, Fulgoromorpha) from mid-Cretaceous amber of northern Myanmar, with new insights in the classification of the family. Cretac. Res. 128, 104959. 10.1016/j.cretres.2021.104959

Maksoud, S., Azar, D., Granier, B., Gèze, R., 2017. New data on the age of the Lower Cretaceous amber outcrops of Lebanon. Palaeoworld 26, 331–338. 10.1016/j.palwor.2016.03.003

Manni, M., Berkeley, M.R., Seppey, M., Simão, F.A., Zdobnov, E.M., 2021. BUSCO Update: Novel and Streamlined Workflows along with Broader and Deeper Phylogenetic Coverage for Scoring of Eukaryotic, Prokaryotic, and Viral Genomes. Mol. Biol. Evol. 38, 4647–4654. 10.1093/molbev/msab199

Martínez-Delclòs, X., Briggs, D.E.G., Peñalver, E., 2004. Taphonomy of insects in carbonates and amber. Palaeogeogr. Palaeoclimatol. Palaeoecol. 203, 19–64. 10.1016/S0031-0182(03)00643-6

Matthews, K.J., Maloney, K.T., Zahirovic, S., Williams, S.E., Seton, M., Müller, R.D., 2016. Global plate boundary evolution and kinematics since the late Paleozoic. Glob. Planet. Change 146, 226–250. 10.1016/j.gloplacha.2016.10.002

McCutcheon, J.P., Boyd, B.M., Dale, C., 2019. The life of an insect endosymbiont from the cradle to the grave. Curr. Biol. 29, R485–R495. 10.1016/j.cub.2019.03.032

Metcalf, Z.P., 1947. Fulgoridae, in: General Catalogue of the Hemiptera. Fascicle IV Fulgoroidea, Part 9. Smith College, Northampton, MA.

Metcalf, Z.P., 1938. The Fulgorina of Barro Colorado and other parts of Panama. Bull. Mus. Comp. Zool. 82, 275–423.

Michalik, A., Franco, D.C., Deng, J., Szklarzewicz, T., Stroiński, A., Kobiałka, M., Łukasik, P., 2023. Variable organization of symbiont-containing tissue across planthoppers hosting different heritable endosymbionts. Front. Physiol. 14, 1135346. 10.3389/fphys.2023.1135346

Minh, B.Q., Schmidt, H.A., Chernomor, O., Schrempf, D., Woodhams, M.D., von Haeseler, A., Lanfear, R., 2020. IQ-TREE 2: New Models and Efficient Methods for Phylogenetic Inference in the Genomic Era. Mol. Biol. Evol. 37, 1530–1534. 10.1093/molbev/msaa015

Misof, B., Liu, S., Meusemann, K., Peters, R.S., Donath, A., Mayer, C., Frandsen, P.B., Ware, J., Flouri, T., Beutel, R.G., Niehuis, O., Petersen, M., Izquierdo-Carrasco, F., Wappler, T., Rust, J., Aberer, A.J., Aspock, U., Aspock, H., Bartel, D., Blanke, A., Berger, S., Bohm, A., Buckley, T.R., Calcott, B., Chen, J., Friedrich, F., Fukui, M., Fujita, M., Greve, C., Grobe, P., Gu, S., Huang, Y., Jermiin, L.S., Kawahara, A.Y., Krogmann, L., Kubiak, M., Lanfear, R., Letsch, H., Li, Y., Li, Z., Li, J., Lu, H., Machida, R., Mashimo, Y., Kapli, P., McKenna, D.D., Meng, G., Nakagaki, Y., Navarrete-Heredia, J.L., Ott, M., Ou, Y., Pass, G., Podsiadlowski, L., Pohl, H., von Reumont, B.M., Schutte, K., Sekiya, K., Shimizu, S., Slipinski, A., Stamatakis, A., Song, W., Su, X., Szucsich, N.U., Tan, M., Tan, X., Tang, M., Tang, J., Timelthaler, G., Tomizuka, S., Trautwein, M., Tong, X., Uchifune, T., Walzl, M.G., Wiegmann, B.M., Wilbrandt, J., Wipfler, B., Wong, T.K.F., Wu, Q., Wu, G., Xie, Y., Yang, S., Yang, Q., Yeates, D.K., Yoshizawa, K., Zhang, Q., Zhang, R., Zhang, W., Zhang, Y., Zhao, J., Zhou, C., Zhou, L., Ziesmann, T., Zou, S., Li, Y., Xu, X., Zhang, Y., Yang, H., Wang, J., Wang, J., Kjer, K.M., Zhou, X., 2014. Phylogenomics resolves the timing and pattern of insect evolution. Science 346, 763–767. 10.1126/science.1257570

Muir, F.A.G., 1923. On the classification of the Fulgoroidea (Homoptera). Proc. Hawaii. Entomol. Soc. 5, 205–247.

O’Brien, L.B., 2002. The Wild Wonderful World of Fulgoromorpha. Denisia 04 Zugleich Kat. OÖ Landesmus. Linz N. F. Nr 176 83–102.

Peris, D., Condamine, F.L., 2024. The angiosperm radiation played a dual role in the diversification of insects and insect pollinators. Nat. Commun. 15, 552. 10.1038/s41467-024-44784-4

Price, M.N., Dehal, P.S., Arkin, A.P., 2010. FastTree 2 – Approximately Maximum-Likelihood Trees for Large Alignments. PLoS ONE 5, e9490. 10.1371/journal.pone.0009490

Prjibelski, A., Antipov, D., Meleshko, D., Lapidus, A., Korobeynikov, A., 2020. Using SPAdes De Novo Assembler. Curr. Protoc. Bioinforma. 70. 10.1002/cpbi.102

Rambaut, A., Drummond, A.J., Xie, D., Baele, G., Suchard, M.A., 2018. Posterior Summarization in Bayesian Phylogenetics Using Tracer 1.7. Syst. Biol. 67, 901–904. 10.1093/sysbio/syy032

Schachat, S.R., Goldstein, P.Z., Desalle, R., Bobo, D.M., Boyce, C.K., Payne, J.L., Labandeira, C.C., 2022. Illusion of flight? Absence, evidence and the age of winged insects. Biol. J. Linn. Soc. 138, 143–168. 10.1093/biolinnean/blac137

Schmidt, E., 1915. Die Dictyopharinen des Stettiner Museums. (Hemiptera-Homoptera). Entomologische Zeitung. Herausgegeben von dem entomologischen Vereine zu Stettin. Stettin 76: 345–358 [358]. Entomol. Ztg. Hrsg. Von Dem Entomol. Vereine Zu Stettin Stettin 76, 345–358.

Simion, P., Delsuc, F., Philippe, H., 2020. To What Extent Current Limits of Phylogenomics Can Be Overcome?, in: Phylogenetics in the Genomic Era. No commercial publisher | Authors open access book, p. 2.1:1--2.1:34.

Simon, C., Gordon, E.R.L., Moulds, M.S., Cole, J.A., Haji, D., Lemmon, A.R., Lemmon, E.M., Kortyna, M., Nazario, K., Wade, E.J., Meister, R.C., Goemans, G., Chiswell, S.M., Pessacq, P., Veloso, C., Mccutcheon, J.P., Łukasik, P., 2019. Off-target capture data, endosymbiont genes and morphology reveal a relict lineage that is sister to all other singing cicadas. Biol. J. Linn. Soc. 128, 865–886.

Skinner, R.K., Dietrich, C.H., Walden, K.K.O., Gordon, E., Sweet, A.D., Podsiadlowski, L., Petersen, M., Simon, C., Takiya, D.M., Johnson, K.P., 2020. Phylogenomics of Auchenorrhyncha (Insecta: Hemiptera) using transcriptomes: examining controversial relationships via degeneracy coding and interrogation of gene conflict. Syst. Entomol. 45, 85–113. 10.1111/syen.12381

Song, N., Liang, A.-P., 2013. A Preliminary Molecular Phylogeny of Planthoppers (Hemiptera: Fulgoroidea) Based on Nuclear and Mitochondrial DNA Sequences. PLoS ONE 8, e58400. 10.1371/journal.pone.0058400

Song, Z.-S., Bartlett, C.R., O’Brien, L.B., Liang, A.-P., Bourgoin, T., 2018. Morphological phylogeny of Dictyopharidae (Hemiptera: Fulgoromorpha). Syst. Entomol. 43, 637–658. 10.1111/syen.12293

Song, Z.-S., Szwedo, J., Wang, R.-R., Liang, A.-P., 2016. Systematic revision of Aluntiini Emeljanov, 1979 (Hemiptera: Fulgoromorpha: Dictyopharidae: Dictyopharinae): reclassification, phylogenetic analysis, and biogeography. Zool. J. Linn. Soc. 176, 349–398. 10.1111/zoj.12319

Stroiński, A., Bourgoin, T., Szwedo, J., 2022. Laberiini, a new tribe of Tropiduchidae planthoppers from Madagascar (Hemiptera: Fulgoroidea). Eur. J. Taxon. 836, 23–54. 10.5852/ejt.2022.836.1913

Szwedo, J., 2017. The unity, diversity and conformity of bugs (Hemiptera) through time. Earth Environ. Sci. Trans. R. Soc. Edinb. 107, 109–128. 10.1017/S175569101700038X

Szwedo, J., Bourgoin, T., Lefebvre, F., 2004. Fossil Planthoppers (Hemiptera: Fulgoromorpha) of the World. An annotated catalogue with notes on Hemiptera classification.

Studio 1, Warszawa. Urban, J.M., Bartlett, C.R., Cryan, J.R., 2010. Evolution of Delphacidae (Hemiptera: Fulgoroidea): combined-evidence phylogenetics reveals importance of grass host shifts. Syst. Entomol. 35, 678–691. 10.1111/j.1365-3113.2010.00539.x

Urban, J.M., Cryan, J.R., 2012. Two ancient bacterial endosymbionts have coevolved with the planthoppers (Insecta: Hemiptera: Fulgoroidea). BMC Evol. Biol. 12, 87. 10.1186/1471-2148-12-87

Urban, J.M., Cryan, J.R., 2009. Entomologically famous, evolutionarily unexplored: The first phylogeny of the lanternfly family Fulgoridae (Insecta: Hemiptera: Fulgoroidea). Mol. Phylogenet. Evol. 50, 471–484. 10.1016/j.ympev.2008.12.004

Urban, J.M., Cryan, J.R., 2007. Evolution of the planthoppers (Insecta: Hemiptera: Fulgoroidea). Mol. Phylogenet. Evol. 42, 556–572. 10.1016/j.ympev.2006.08.009

Viegas, E.F.G., 2023. Sistematica e biogeografia de Achilixiidae Muir (Hemiptera: Auchenorrhyncha: Fulgoroidea) com base em caracteres morfológicos e molecularesTese. Tese Inst. Nac. Pesqui. Amaz. INPA Manaus Braz. 1–230.

Viegas, E.F.G., Ale-Rocha, R., 2024. A century of Achilixiidae Muir, 1923 (Hemiptera: Auchenorrhyncha: Fulgoromorpha): taxonomic study of the genus Bebaiotes Muir, 1924 and description of eight new species from Brazil. Zootaxa 5413, 1–65. 10.11646/zootaxa.5413.1.1

Wang, L.-G., Lam, T.T.-Y., Xu, S., Dai, Z., Zhou, L., Feng, T., Guo, P., Dunn, C.W., Jones, B.R., Bradley, T., Zhu, H., Guan, Y., Jiang, Y., Yu, G., 2019. Treeio: An R Package for Phylogenetic Tree Input and Output with Richly Annotated and Associated Data. Mol. Biol. Evol. 37, 599–603. 10.1093/molbev/msz240

Wang, M., Zhang, Y., Bourgoin, T., 2016. Planthopper family Issidae (Insecta: Hemiptera: Fulgoromorpha): linking molecular phylogeny with classification. Mol. Phylogenet. Evol. 105, 224–234. 10.1016/j.ympev.2016.08.012

Wang, R.-R., Li, X.-Y., Szwedo, J., Stroiński, A., Liang, A.-P., Bourgoin, T., 2017. Testing Tropiduchini Stål 1866 (Hemiptera: Tropiduchidae) monophyly, a young inter-tropical taxon of mainly insular species: taxonomy, distribution patterns and phylogeny, with the description of a new genus from Papua New Guinea. Syst. Entomol. 42, 359–378. 10.1111/syen.12219

Wang, R.-R., Stroiński, A., Szwedo, J., Bourgoin, T., Liang, A.-P., 2014. Recent Dispersal and Diet Relaxation Might Explain the Monotypic and Endemic Genus Montrouzierana Signoret, 1861 in New Caledonia (Hemiptera: Fulgoromorpha: Tropiduchidae). Ann. Zool. 64, 693–708. 10.3161/000345414X685974

Wang, W., Meng, R., Huang, Y., Fang, W., Zhang, H., Liu, H., Stroiński, A., Bourgoin, T., Qin, D., 2023. A phylogeny with divergence-time estimation of planthoppers (Hemiptera: Fulgoroidea) based on mitochondrial sequences. Zool. J. Linn. Soc. zlad110, 1–12. 10.1093/zoolinnean/zlad110

Wheeler, T.J., Eddy, S.R., 2013. nhmmer: DNA homology search with profile HMMs. Bioinformatics 29, 2487–2489. 10.1093/bioinformatics/btt403

Wilson, S.W., Wheeler, A.G., 1992. Host Plant and Descriptions of Nymphs of the Planthopper *Rhabdocephala brunnea* (Homoptera: Fulgoridae). Ann. Entomol. Soc. Am. 85, 258–264. 10.1093/aesa/85.3.258

Yang, L.-F., Hu, C.-L., 1985. Description of four new species of Meenoplidae (Homoptera: Fulgoroidea). Nanjing Nongye Daxue Xuebao J Nanjing Agric Univ 21–27.

Yeh, W.-B., Yang, C.-T., Hui, C.-F., 2005. A Molecular Phylogeny of Planthoppers (Hemiptera: Fulgoroidea) Inferred from Mitochondrial 16S rDNA Sequences. Zool. Stud. 44, 519–535.

Zhang, C., 2022. Weighting by Gene Tree Uncertainty Improves Accuracy of Quartet-based Species Trees. Mol. Biol. Evol. 39, msac215. 10.1093/molbev/msac215

Zhang, C., 2018. ASTRAL-III: polynomial time species tree reconstruction from partially resolved gene trees. BMC Bioinformatics 19, 153. 10.1186/s12859-018-2129-y

Zuntini, A.R., Carruthers, T., Maurin, O., Bailey, P.C., Leempoel, K., 2024. Phylogenomics and the rise of the angiosperms. Nature. 10.1038/s41586-024-07324-0

