## Supplementary material for "Phylogenomics resolves the relationship and the evolutionary history of planthoppers (Insecta: Hemiptera: Fulgoromorpha)": Figures. S1-8

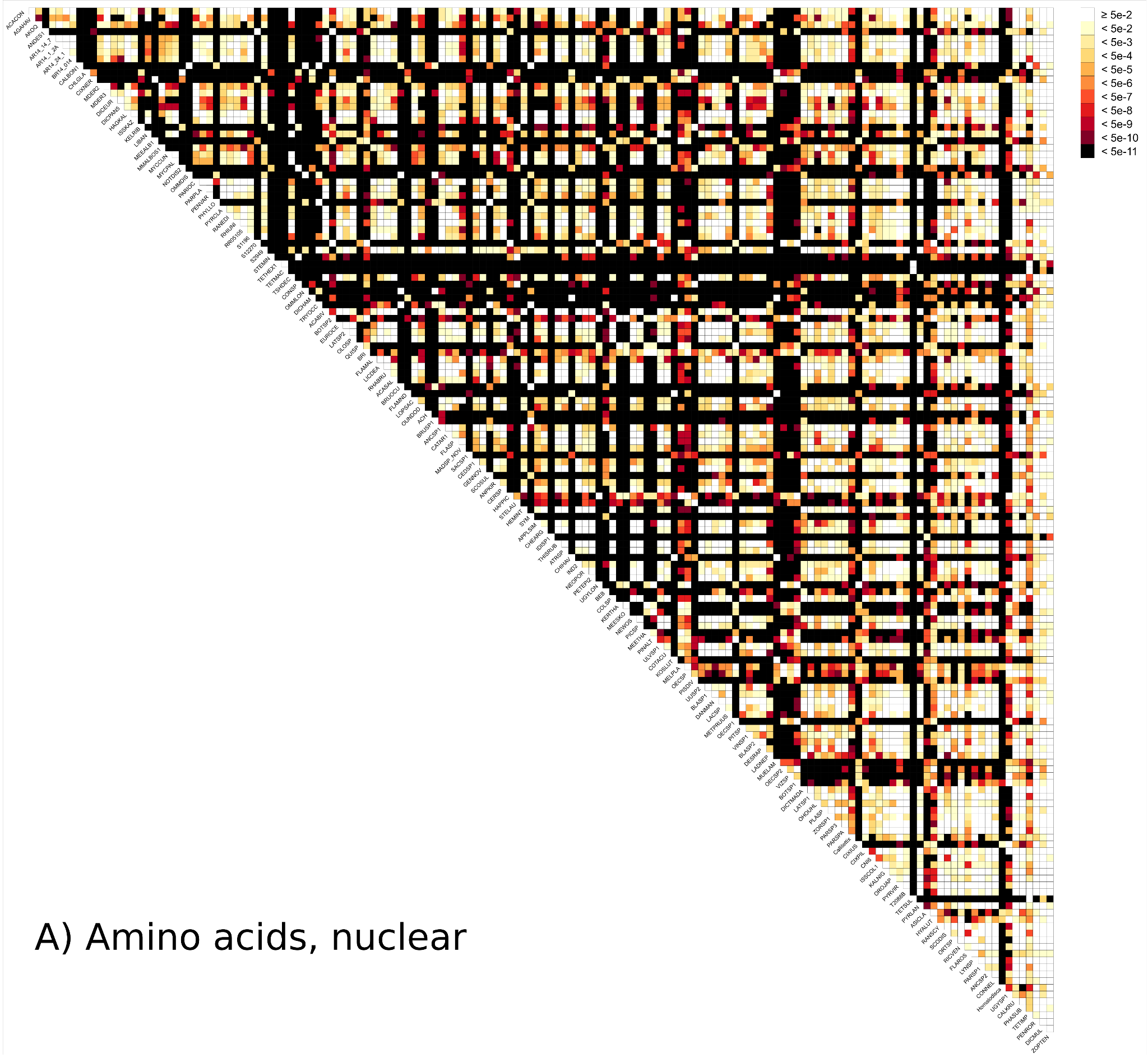

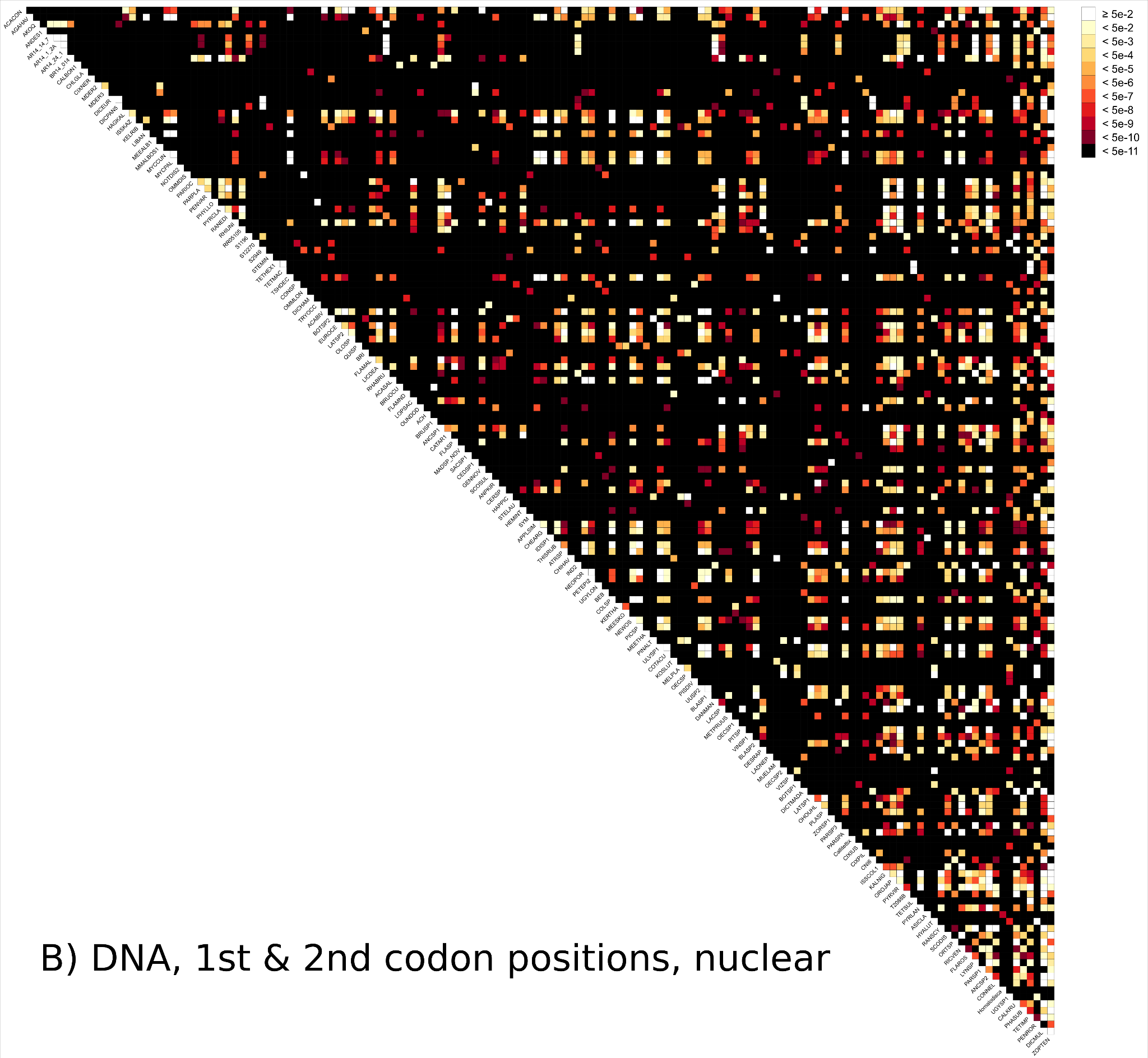

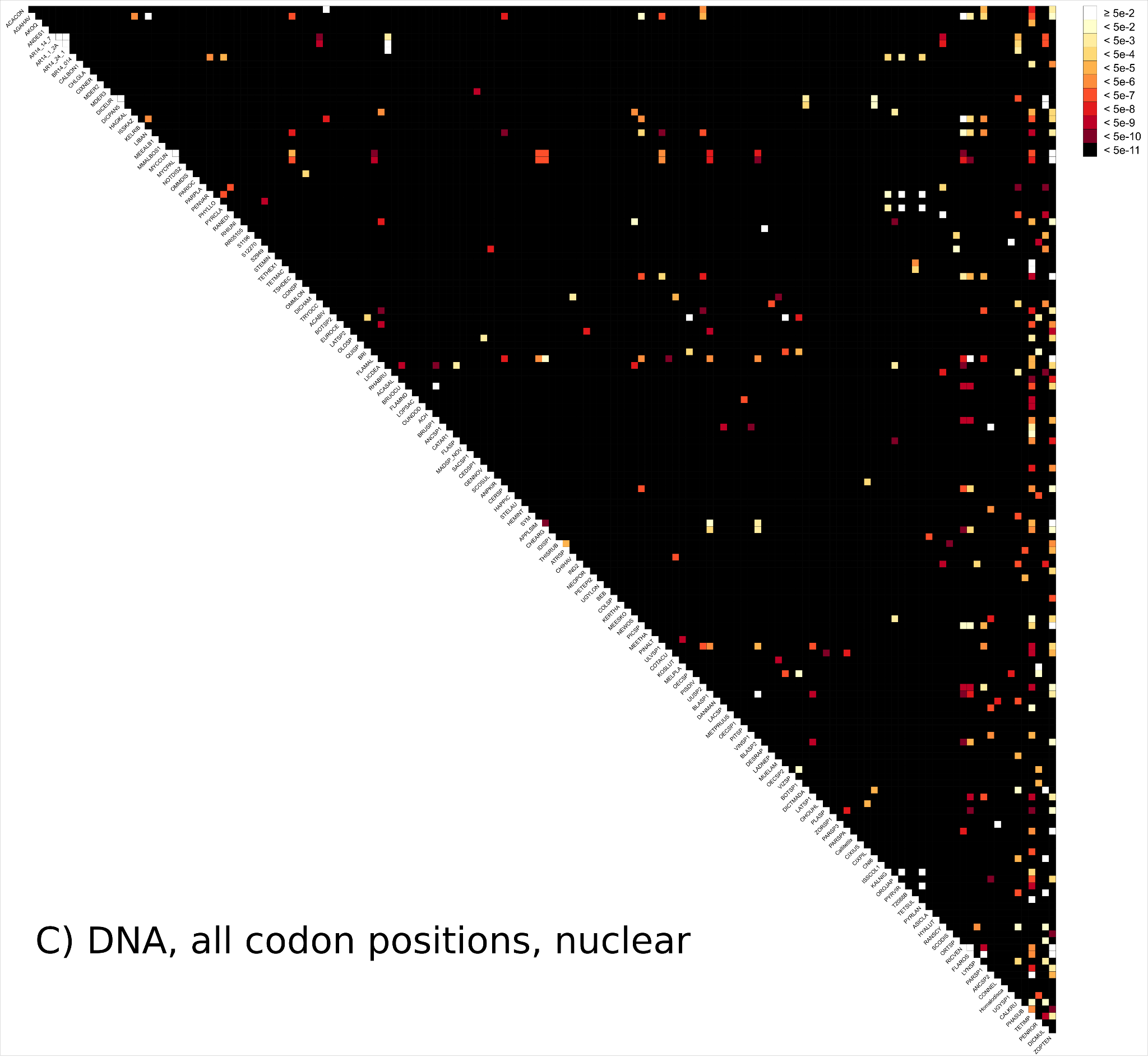

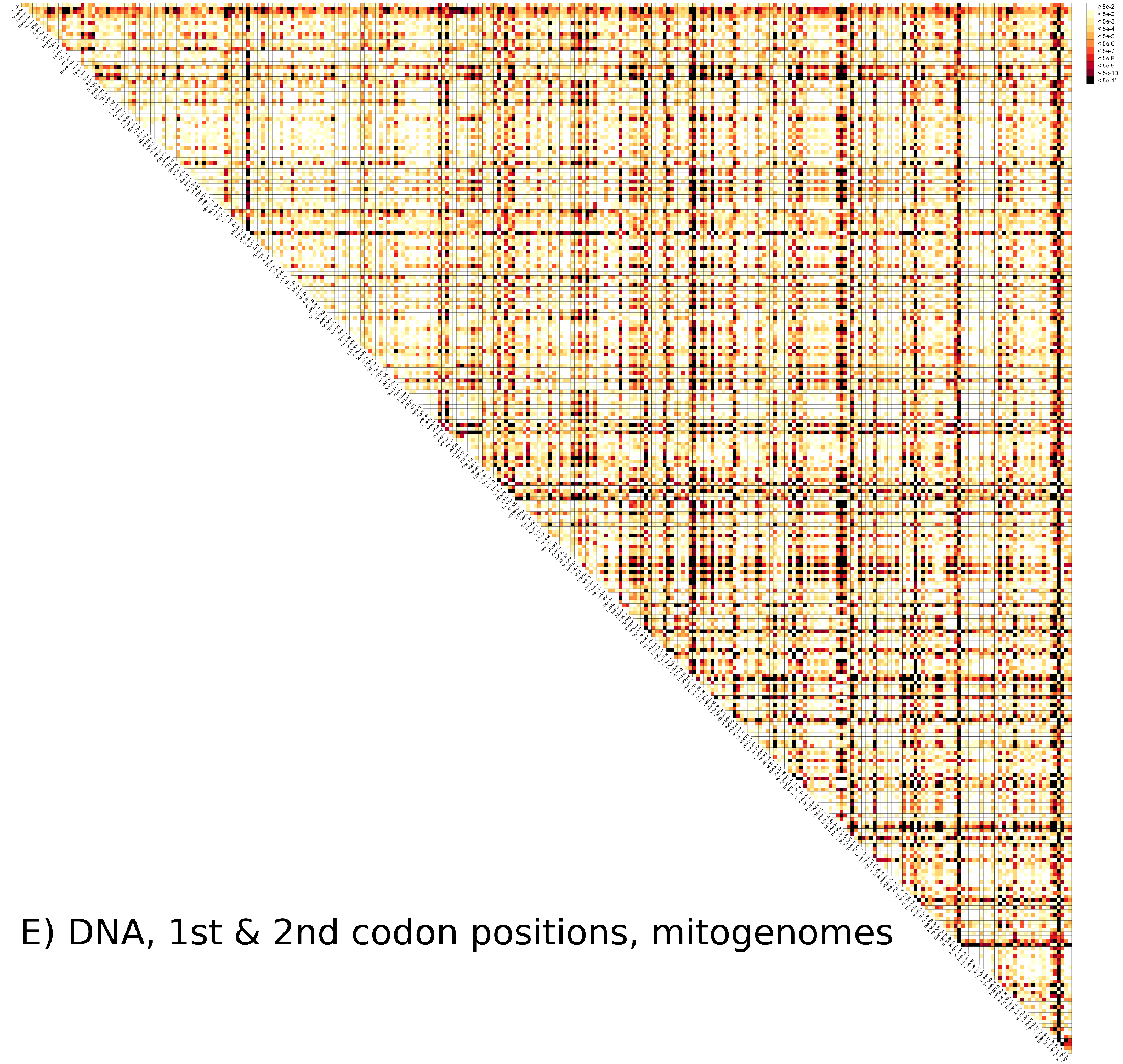

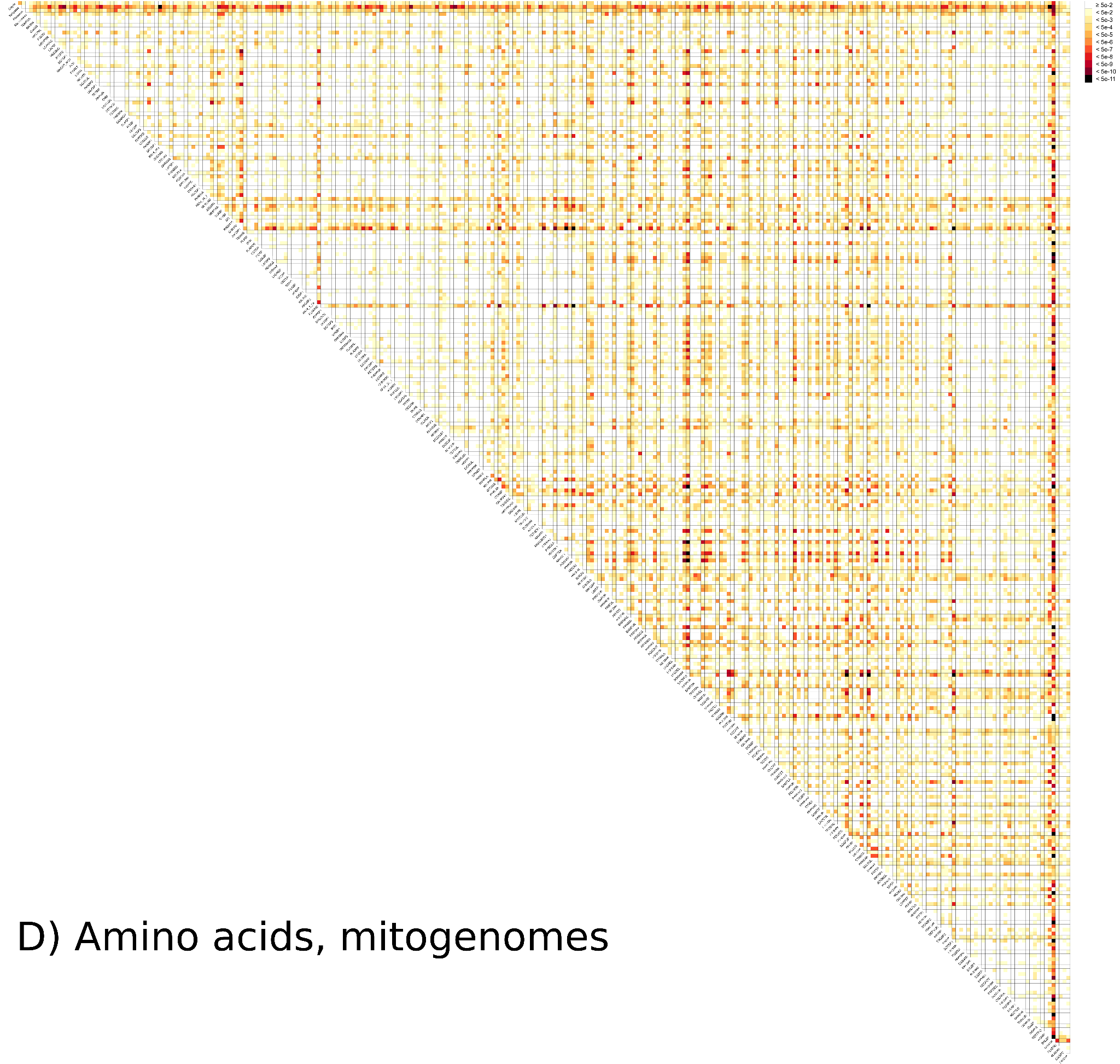
**Figure. S1**
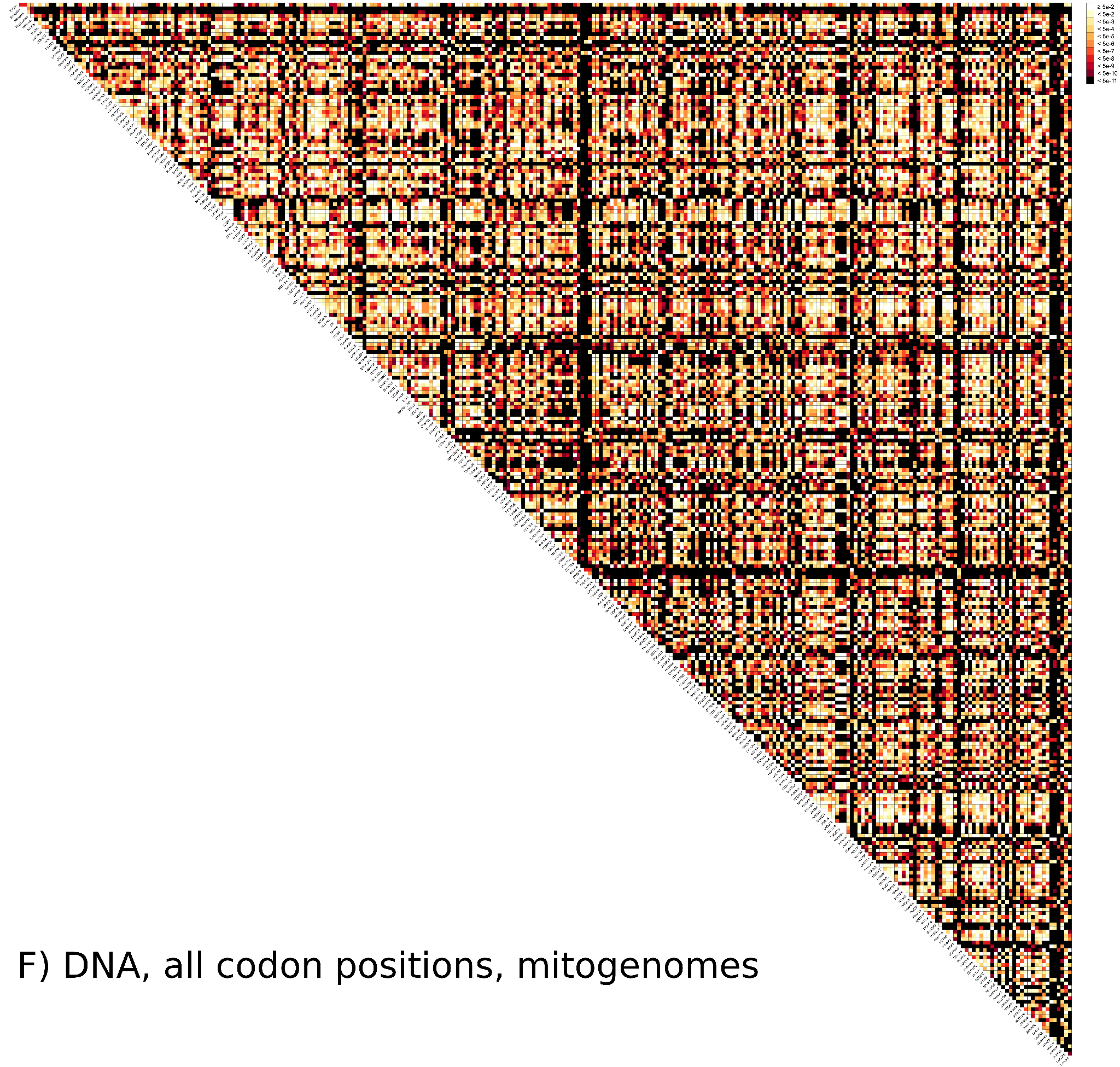
 Test for among lineage compositional heterogeneity. Heat map showing pairwise Bowker's tests of A) D) the amino acid supermatrix, B) E) the nucleotide supermatrix including only 1st and 2nd codon positions, and C) F) the nucleotide supermatrix including all codon positions of nuclear and mitochondrial datasets, respectively. P-values > 0.05 are colored in white and indicate sequence pairs that fully match globally stationary, reversible, and homogeneous (SRH) conditions.

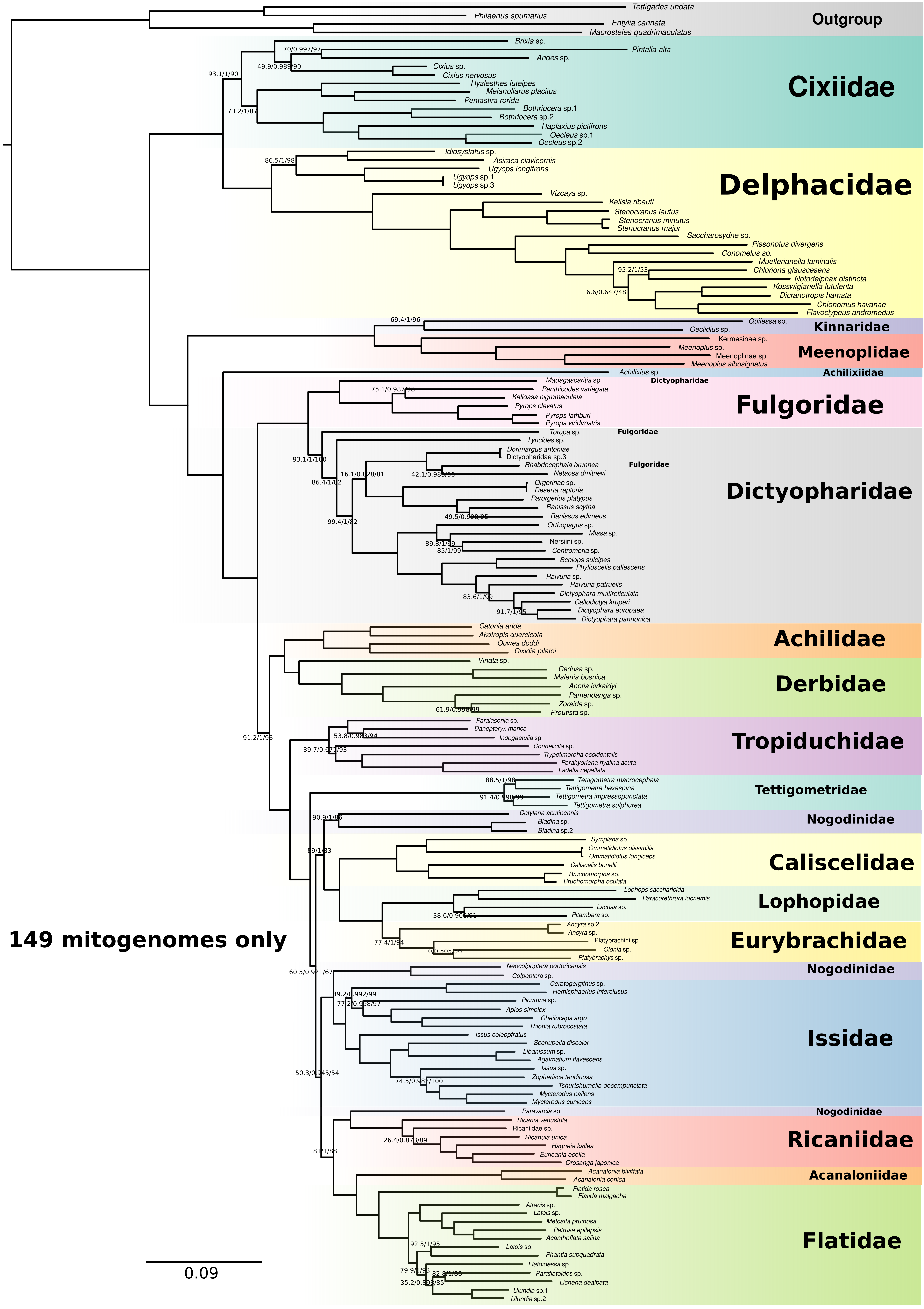

**Figure S2**. The best ML tree based on the concatenated nucleotide dataset of 13 mitochondrial protein coding genes (PCGs) of 149 planthopper species newly sequenced in this study. The branch supports that either the SH-aLRT (left) or the ultrafast bootstrap support (right) is less than 95 are shown, with the approximate Bayes test bootstrap in the middle.

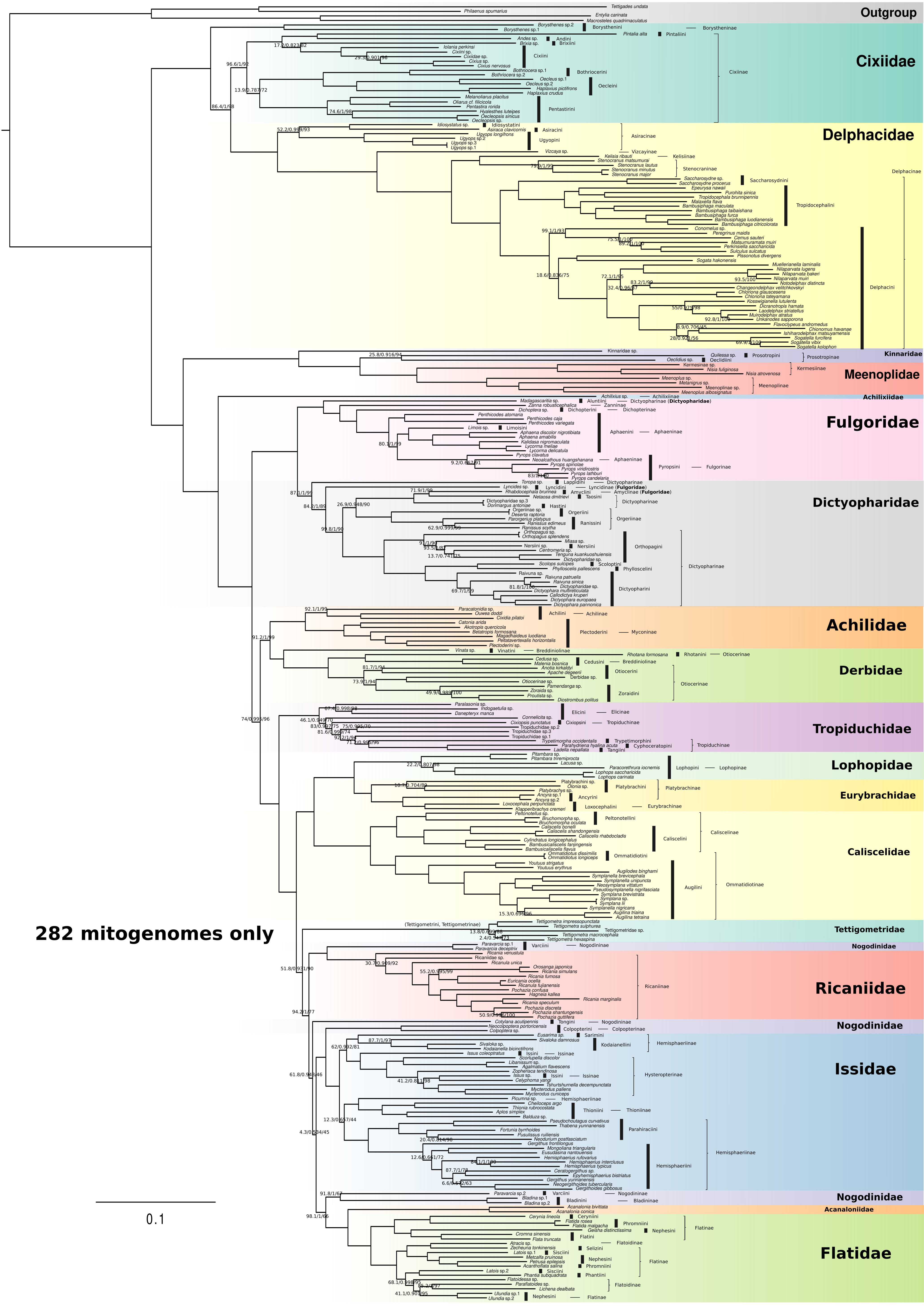

**Figure S3**. The best ML tree based on the concatenated nucleotide dataset of 13 mitochondrial PCGs of 282 planthopper species. The branch supports that either the SH-aLRT (left) or the ultrafast bootstrap support (right) is less than 95 are shown, with the approximate Bayes test bootstrap in the middle. The tribe and subfamily information are indicated next to each species according to Fulgoromorpha Lists On the Web (FLOW; flow.hemiptera-databases.org/flow/).

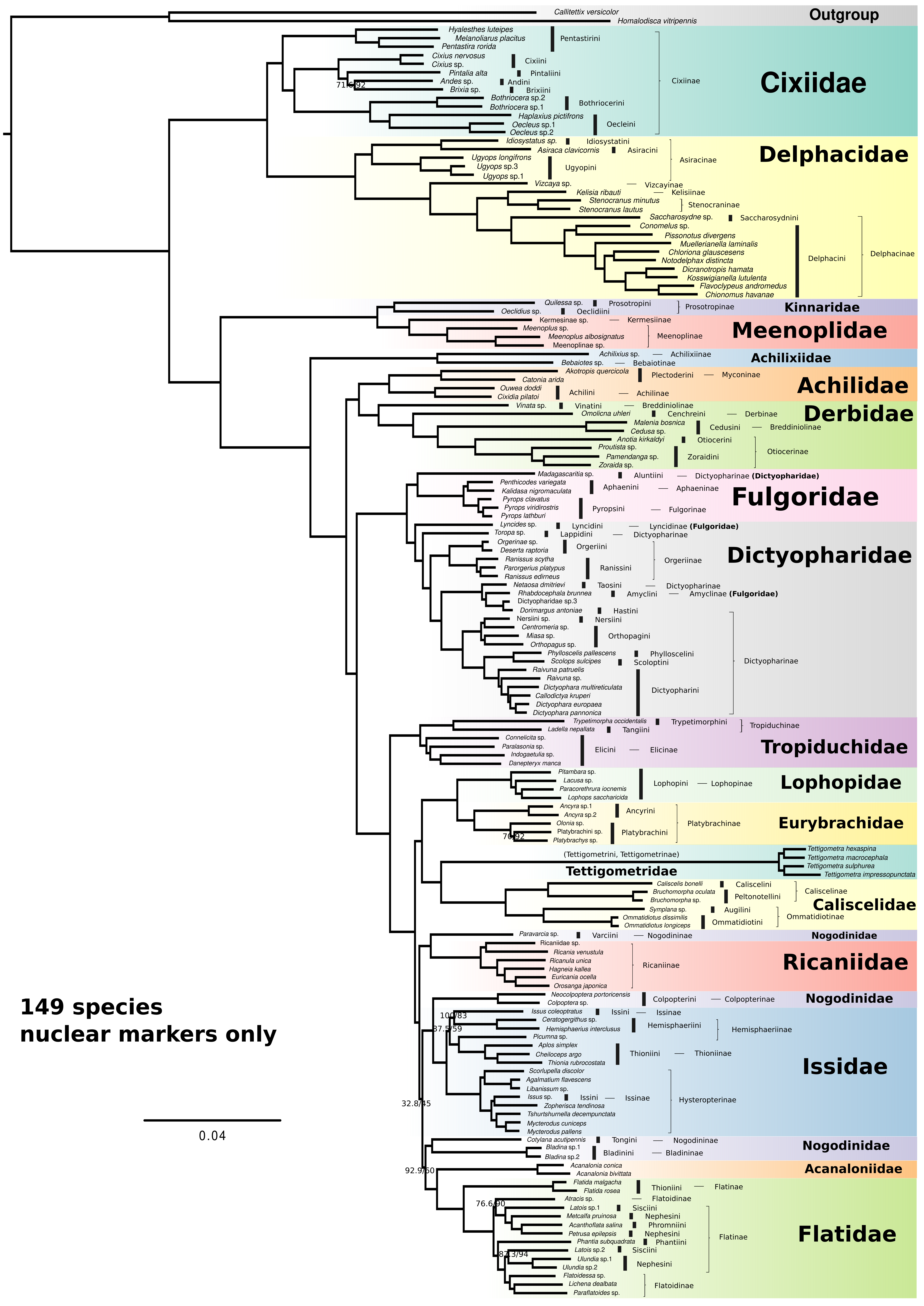

**Figure S4**. The best ML tree based on the concatenated nucleotide dataset of 1164 nuclear markers of 149 planthopper species newly sequenced in this study. The branch supports that either the SH-aLRT (left) or the ultrafast bootstrap support (right) is less than 95 are shown. The approximate Bayes test gives full support to all branches and therefore is not shown. The tribe and subfamily information are indicated next to each species according to Fulgoromorpha Lists On the Web (FLOW; flow.hemiptera-databases.org/flow/).

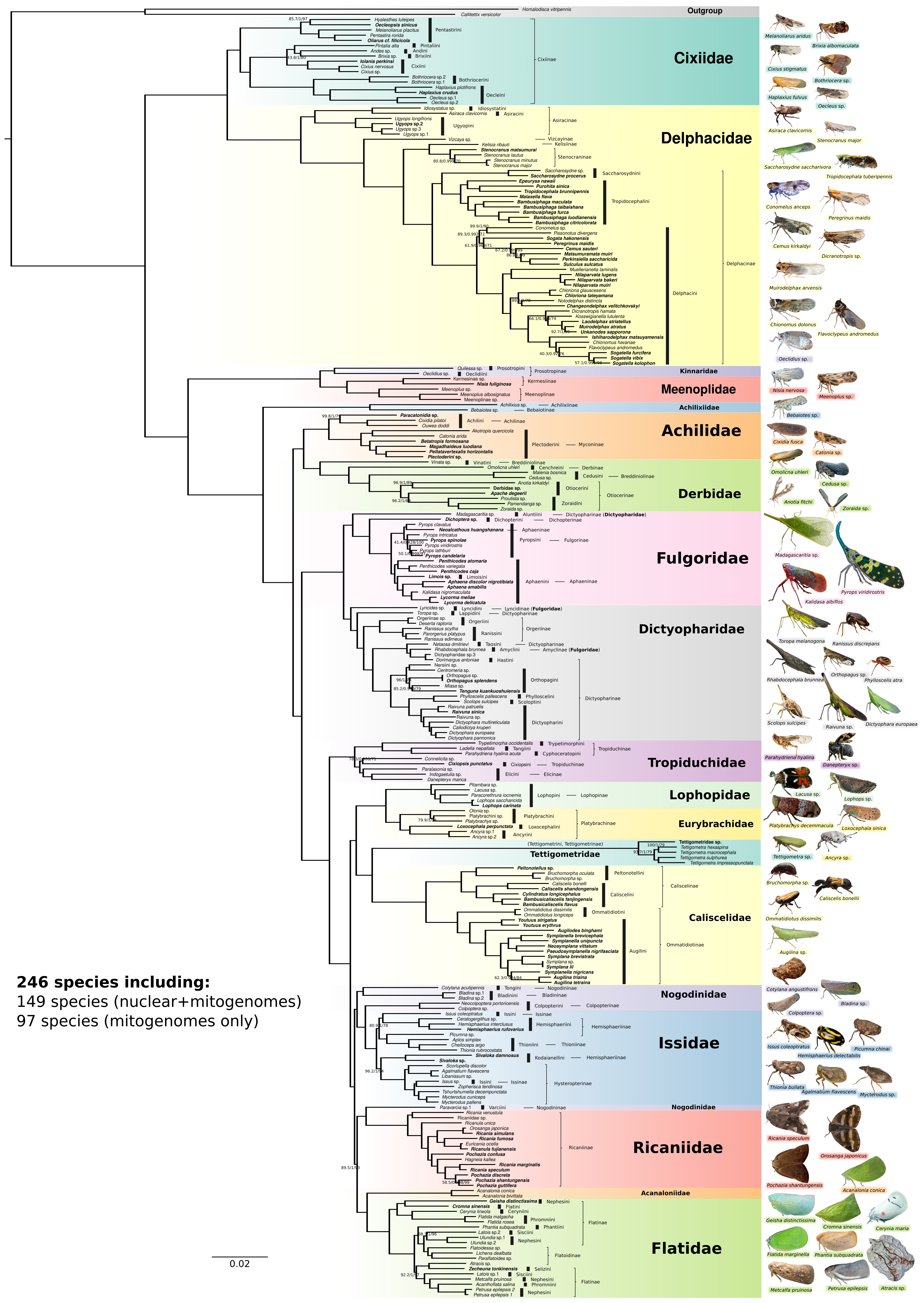

**Figure S5**. The best ML tree based on the concatenated nucleotide dataset of 1164 nuclear markers and 13 mitochondrial PCGs of 246 planthopper species. Forty species from Wang et al. (2023) are excluded from this phylogeny. The branch supports that either the SH-aLRT (left) or the ultrafast bootstrap support (right) is less than 95 are shown, with the approximate Bayes test bootstrap in the middle. Species contributing only mitogenomes have their names in bold. The tribe and subfamily information are indicated next to each species according to Fulgoromorpha Lists On the Web (FLOW; [flow.hemiptera-databases.org/flow/](http://flow.hemiptera-databases.org/flow/)).

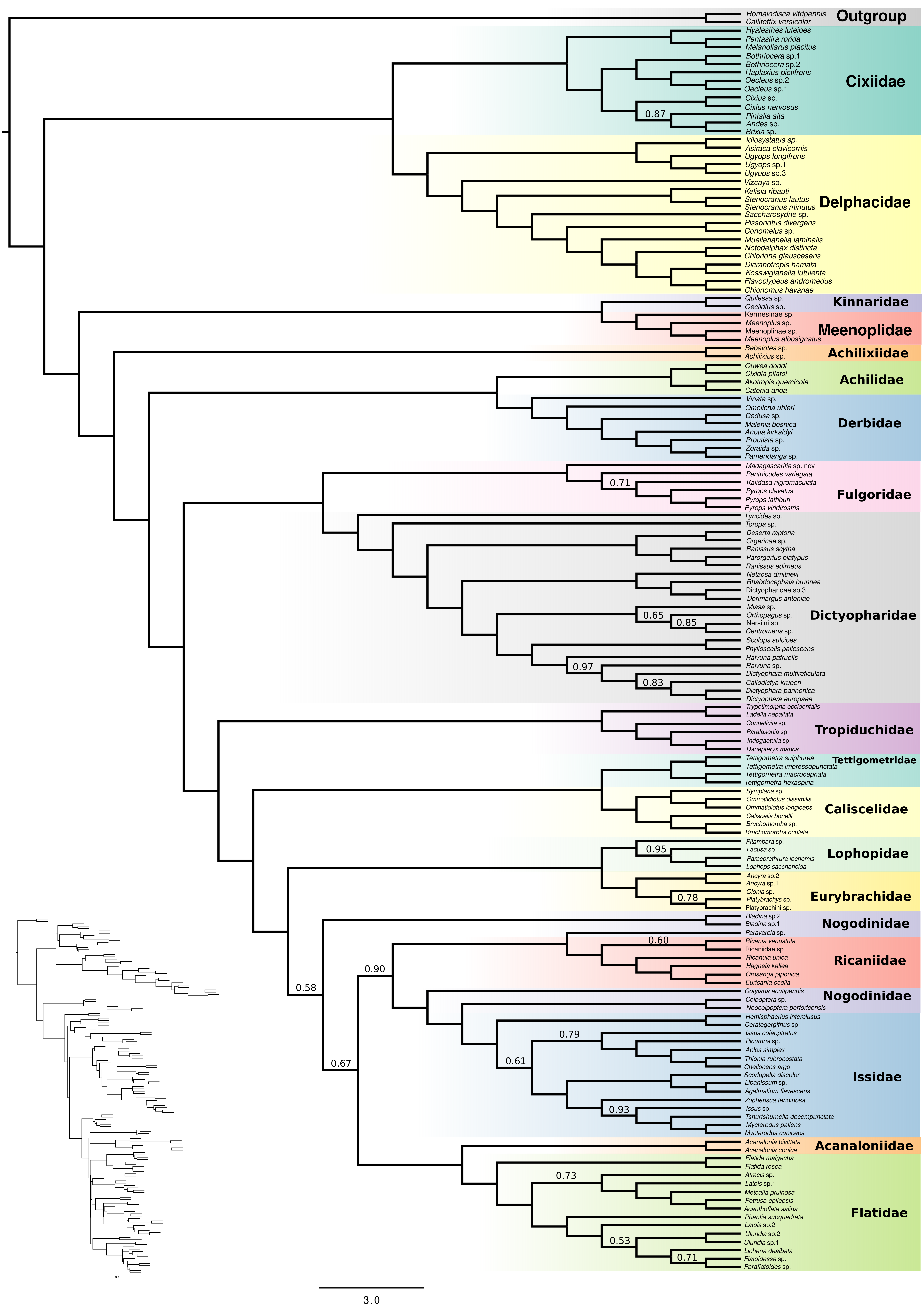

**Figure S6**. The species tree of 149 species inferred from 1164 nuclear gene trees by ASTRAL (Down left). The cladogram is shown for illustration purposes. Branch lengths are shown in coalescent units. Branch supports are shown as local posterior probabilities, which were computed based on gene tree quartet frequencies.

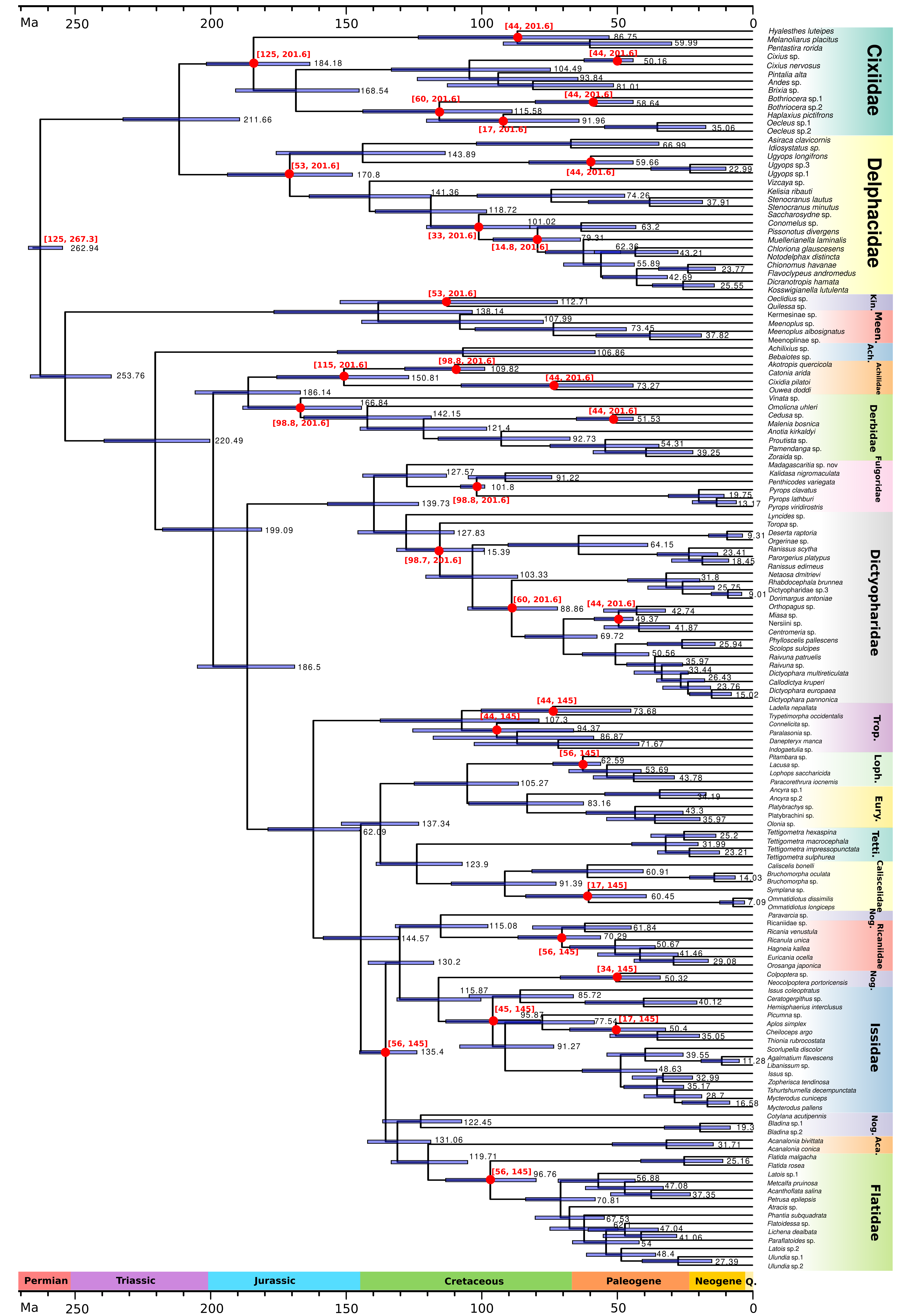

**Figure S7**. The dated phylogeny of 149 planthopper species estimated in BEAST based on a reduced dataset of 57 nuclear genes and the topology of the best ML tree from IQ-TREE2 in Figure S4. All 30 calibration points with fossils are indicated by red circles and were assigned with uniform priors and our age constraints based on fossil history and host plants. The minimum and maximum age constraints of each calibration point are shown in red above or below the node. The maximum constraint for the root is 267.3 Ma. The 95% confidence intervals of node age are noted as light blue bars. The mean ages are shown on the right side of each node bar. Timescales are in millions of years. Family names are abbreviated as follows:: Kinnaridae (Kin.), Meenoplidae (Meen.), Achilixiidae (Ach.), Tropiduchidae (Trop.), Lophopidae (Loph.), Eurybrachidae (Eury.), Tettigometridae (Tetti.), Nogodinidae (Nog.), and Acanaloniidae (Aca.).

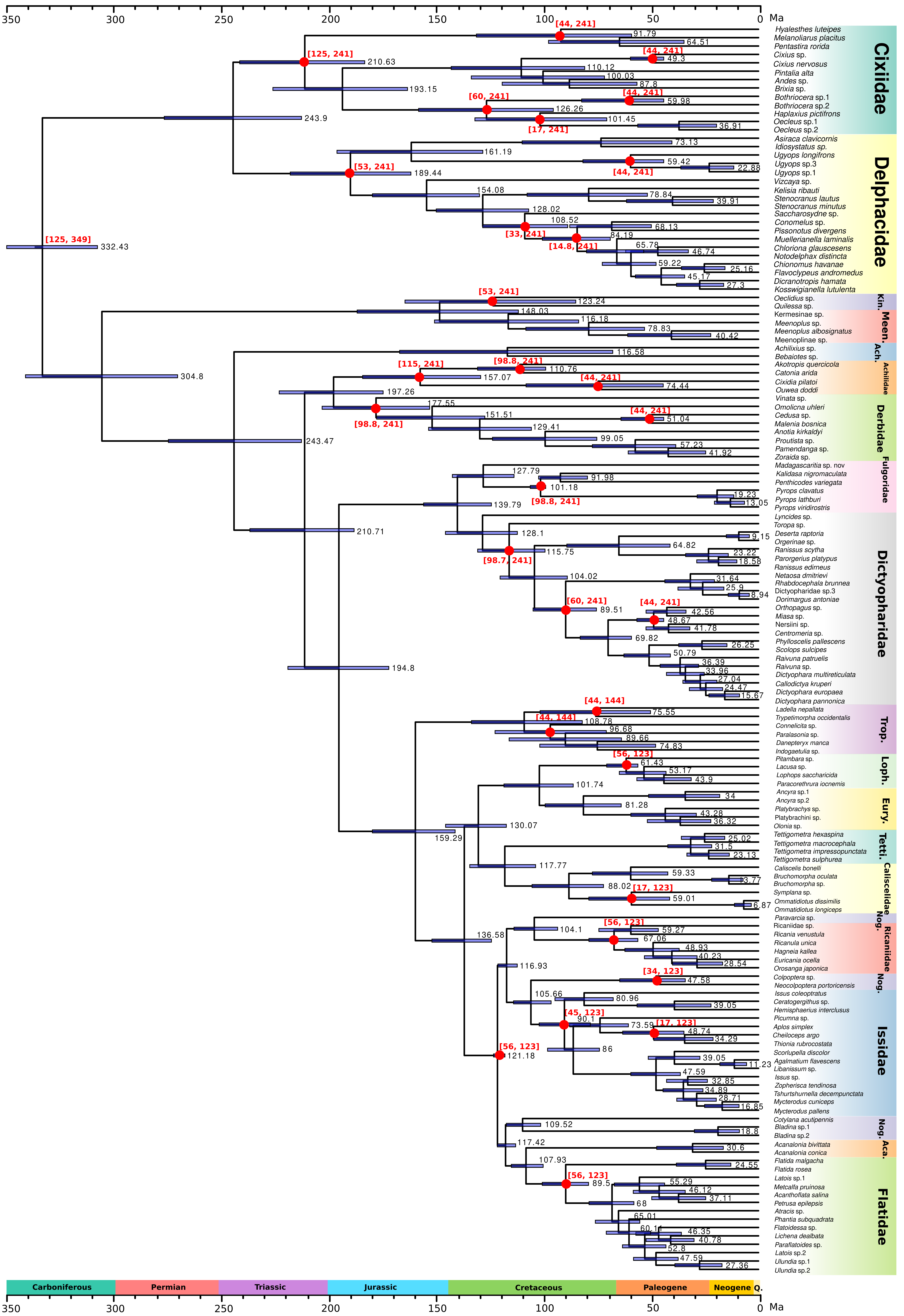

**Figure S8**. A similar dated phylogeny to the one in Figure S7 under the alternative set of age constraints based on the published dated tree from Johnson et al. (2018).
